# An enhancer RNA recruits MLL1 to regulate transcription of *Myb*

**DOI:** 10.1101/2023.09.26.559528

**Authors:** Juhyun Kim, Luis F. Diaz, Matthew J. Miller, Benjamin Leadem, Ivan Krivega, Ann Dean

## Abstract

The *Myb* proto-oncogene encodes the transcription factor c-MYB, which is critical for hematopoiesis. Distant enhancers of *Myb* form a hub of interactions with the *Myb* promoter. We identified a long non-coding RNA (*Myrlin*) originating from the −81 kb murine *Myb* enhancer. *Myrlin* and *Myb* are coordinately regulated during erythroid differentiation. *Myrlin* TSS deletion using CRISPR/Cas9 reduced *Myrlin* and *Myb* expression and LDB1 complex occupancy at the *Myb* enhancers, compromising enhancer contacts and reducing RNA Pol II occupancy in the locus. In contrast, CRISPRi silencing of *Myrlin* left LDB1 and the *Myb* enhancer hub unperturbed, although *Myrlin* and *Myb* expression were downregulated, decoupling transcription and chromatin looping. *Myrlin* interacts with the MLL1 complex. *Myrlin* CRISPRi compromised MLL1 occupancy in the *Myb* locus, decreasing CDK9 and RNA Pol II binding and resulting in Pol II pausing in the *Myb* first exon/intron. Thus, *Myrlin* directly participates in activating *Myb* transcription by recruiting MLL1.

## INTRODUCTION

While the vast majority of the mammalian genome is transcribed, only a small fraction of these transcripts encodes proteins. The functional relevance of most non-coding transcription remains largely unknown. RNA polymerase II (Pol II) non-coding transcripts that are greater than 200 nt in length and lack coding potential are known as long non-coding RNAs (lncRNAs). LncRNAs, much like their protein coding counterparts, can be spliced and polyadenylated, but they are biased towards two-exon transcripts that remain localized to the nucleus ^1–3^. Nuclear-localized lncRNAs can be involved in gene regulation via interactions with chromatin remodelers, histone modifying complexes and transcription factors ^4–8^. LncRNAs derived from enhancer regulatory regions, known as enhancer RNAs or eRNAs, have the potential ability to function together with the enhancers from which they are derived to regulate expression of target genes ^9–11^.

Enhancers increase the transcriptional output of target genes in a cell-type specific fashion, often bridging substantial genomic distances ^12–17^. Both active enhancers and genes are marked by histone H3 lysine 4 (H3lys4) methylation. This modification is deposited by the MLL/COMPASS family of histone methyltransferases including SET1A/B and four mixed lineage leukemia complexes (MLL1-4) ^18,19^. MLL1 forms a large macromolecular complex with WDR5, Menin, RbBP5, ASH2L and DPY-30 and interacts with the basic transcription machinery, including RNA Pol II ^20^. MLL1 selectively binds non-methyl CpG DNA through its CXXC domain ^21^. Additionally, MLL1 adaptor subunit WDR5 promotes the recruitment of MLL1 to genomic targets at a subset of genes ^22^. The MLL1 complex is recruited by the lncRNA HOTTIP through direct binding with WDR5, establishing H3K4me3 deposition and driving HOXA gene transcription, while *HoxBlinc* recruits both Set1 and MLL complexes to *hoxb* genes through the SET domain. ^23,24^.

The *Myb* proto-oncogene encodes c-MYB (hereafter MYB), a critical hematopoietic regulator of cell proliferation and differentiation ^25^. MYB is a repressor of human fetal hemoglobin production ^26^. Given that elevation of fetal hemoglobin in adults moderates the symptoms of Sickle Cell Disease and β-Thalassemia, MYB is a potential target of therapeutic manipulation. *Myb* is regulated by microRNAs and by a series of enhancers distributed over more than 100 kb between *Myb* and the adjacent upstream *Hbs1l* gene in mouse and human ^27–30^. In the mouse, five enhancers, −36, −61, −68, −81 and −109 kb, with respect to the *Myb* transcription start site (TSS), establish proximity with the *Myb* promoter in an active chromatin hub ^28^. Repression of *Myb*, which is required for terminal differentiation of erythroid cells, is accompanied by loss of these contacts. The enhancers are occupied by the LDB1 transcription factor complex, which mediates enhancer looping, and reduction of LDB1 using an shRNA compromises formation of the *Myb* enhancer hub ^31,32^. However, how chromatin looping and transcription activation at the *Myb* locus are regulated remains unknown.

Long non-coding RNAs have been linked to erythropoiesis and the regulation of numerous erythroid genes, including the adult β-globin and fetal γ-globin genes ^33–38^. We identified a novel lncRNA derived from the murine −81 kb *Myb* enhancer termed *Myrlin* for *<u>My</u>b* enhancer long <u>in</u>tergenic non-coding RNA. The *Myrlin* transcript was not required for formation of the *Myb* enhancer hub. However, *Myrlin* loss reduced MLL1/WDR5 recruitment in the *Myb* locus and decreased CDK9 and RNA Pol II occupancy. Furthermore, *Myrlin* loss resulted in pausing of RNA Pol II within the *Myb* first exon/intron. These results tie the *Myb* locus lncRNA *Myrlin* to the detailed mechanism of *Myb* transcription activation and suggest novel avenues that could become therapeutic targets for increasing HbF in β-globin hemoglobinopathies.

## RESULTS

### The −81 kb murine *Myb* enhancer contains the TSS for a spliced, long intergenic non-coding RNA

The murine *Myb-Hbs1l* intergenic region contains previously characterized regulatory enhancers ^28^. In MEL cells and in primary erythroid cells, the five enhancers, located −36, −61, −68, −81 and −109 kb upstream of the *Myb* promoter, are occupied by the LDB1 complex that includes DNA binding transcription factors GATA1 and TAL1, bridging protein LMO2 and looping protein LDB1 (Figure 1A) ^28^. ChIP-seq and RNA-seq data for uninduced MEL cells indicates that RNA Pol II occupies several of the enhancer sites, but active RNA transcription is only observed at the −81 kb enhancer (Figure 1A) ^28^.

**Figure 1.**
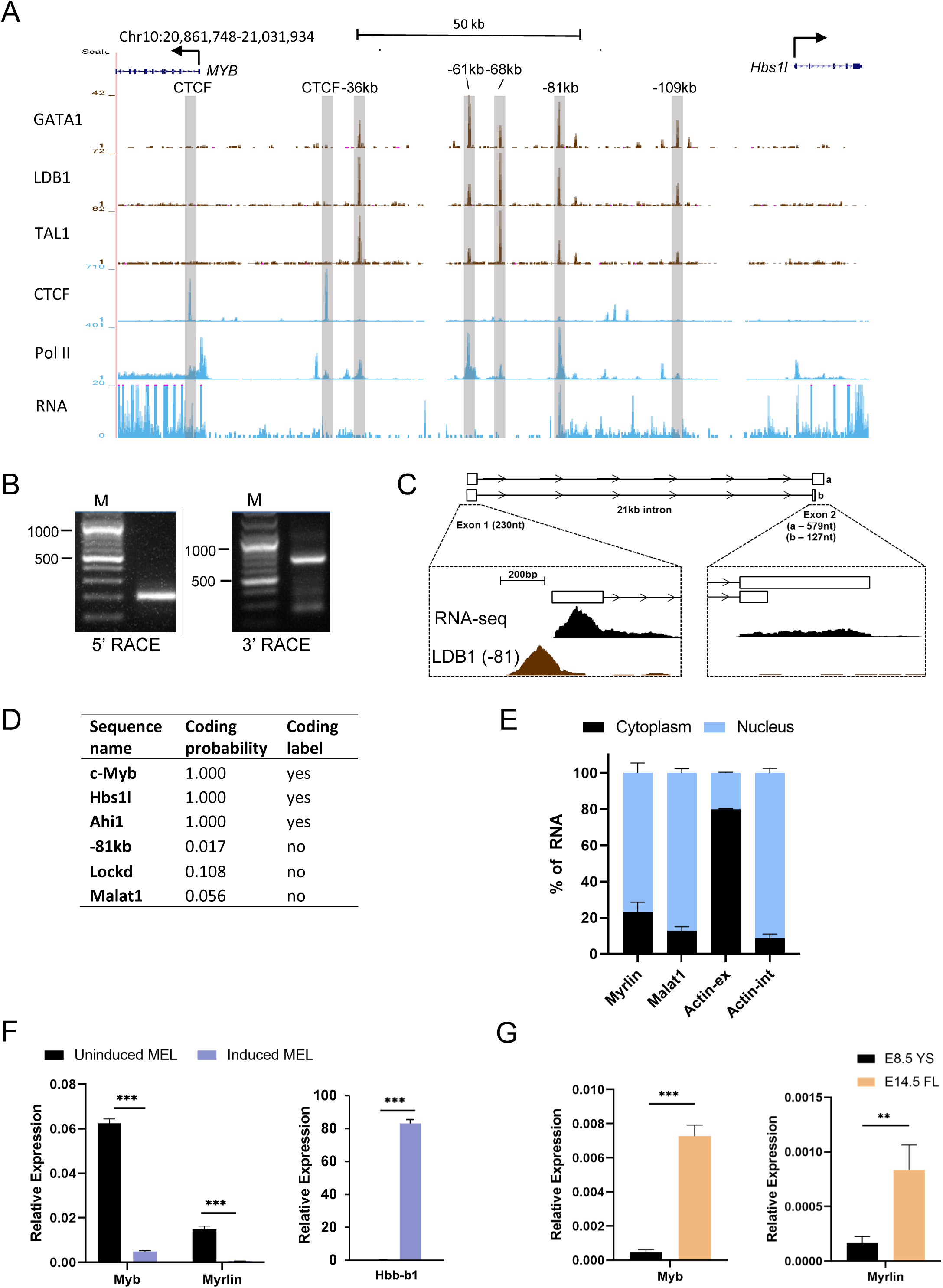
The *Myb* −81 kb enhancer is the transcription start site for the noncoding *Myrlin* RNA. (A) ENCODE ChIP-Seq of LDB1 complex components (GATA1, LDB1 and TAL1), CTCF, and PolII in the *Myb*-*Hbs1l* locus in MEL cells. PolyA RNA-seq is shown in MEL cells. Intergenic LDB1 complex binding sites are highlighted (grey vertical bars). (B) Nested PCR products from 5’ and 3’ RACE in MEL cells. (C) Poly-A RNA-seq and Ldb1 ChIP-seq tracks from ENCODE for MEL at the −81 kb enhancer and downstream *Myrlin* exon 2. (D) Prediction of coding potential for *Myrlin* and select other transcripts as determined by CPAT (see text). (E) Relative expression of *Myrlin* in nuclear and cytoplasmic fractions of MEL cells determined by RT-PCR. *MALAT1* lncRNA and *ActB* provided nuclear and cytoplasmic controls, respectively. (F) Total RNA of uninduced and induced MEL cells was used to determine relative expression of *Myrlin* and *Myb* by RT-PCR. Expression was normalized to *ActB*. (G) Total RNA of E8.5 yolk sac cells (YS) and E14.5 fetal liver cells (FL) was used to determine relative expression of *Myrlin*, *Myb* and *Hbb-b1* by RT-PCR. Expression was normalized to *ActB*. Error bars indicate SEM of 3 independent biological experiments. (*) P < 0.05, (**) P<0.01, (***) P<0.001 by Student’s t-test. See also Figures S1.

Rapid amplification of cDNA ends (5’ and 3’ RACE) revealed an unannotated 2-exon transcript at the *Myb* −81 kb enhancer that exists as two spliced isoforms with a single intron spanning more than 20 kb (Figure 1B, C). The primary transcript is 809 nt with a minor 357 nt shorter form attributable to early termination in exon 2. Stranded RNA-seq from induced and uninduced MEL cells publicly sourced from ENCODE indicates that *Myb* and the −81 kb transcript are divergently transcribed from opposite DNA strands (Figure S1). The transcript has very low coding potential according to the Coding Potential Assessment Tool (CPAT) ^39^ (Figure 1D). Thus, the transcript qualifies as a long non-coding RNA and its low abundance (about 10-fold lower than *Myb*) is consistent with that of an enhancer RNA ^40^.

We named the transcript *Myrlin* (*<u>My</u>b* enhance<u>r l</u>ong <u>in</u>tergenic non-coding RNA). In uninduced MEL cells *Myrlin* is primarily nuclear localized, consistent with a potential role in gene regulation (Figure 1E). Expression of *Myb* decreases upon erythroid cell maturation, which is mirrored by decreases in both *Myb* and *Myrlin* upon differentiation of MEL cells by DMSO (Figure 1F). Like *Myb*, *Myrlin* is expressed at very low levels in E8.5 yolk sac primitive erythroid cells and then more robustly in E14.5 fetal liver definitive erythroid cells ^41^ (Figure 1G). These findings show that expression of *Myb* and *Myrlin* is coordinately regulated in a developmental stage-specific fashion in murine erythroid cells and raises the possibility that *Myrlin* may play a role in regulating *Myb* expression.

### Deletion of the *Myrlin* TSS reduces *Myb* expression

JASPAR motif analysis (http://jaspar.binf.ku.dk/) ^42^ identified a TATA box located −25 nucleotides upstream of the 5’ end of *Myrlin* as determined by RACE, and a GATA1 binding motif, site of LDB1 complex occupancy, located −56 nucleotides upstream (Figure 2A, Figure S2). This organization is consistent with the finding that most mouse erythroid-expressed non-coding RNAs are transcribed from conventional promoters regulated by known transcription factors and are regulated by similar Pol II release mechanisms ^35,43^.To investigate a role for *Myrlin* in *Myb* expression, we generated several MEL cell lines with small deletions that were designed to target the transcription start site (TSS) of *Myrlin* (Figure 2A, Figure S2). Three different mutations were obtained all of which reduced *Myrlin* and *Myb* transcription to varying degrees in uninduced MEL cells (Figure 2B). We chose for further study the 17 base pair deletion (Δ17, hereafter ΔTSS), which removes most of the sequence between the TATA box and the initiator element, and results in the strongest reduction of *Myrlin* and *Myb*.

**Figure 2.**
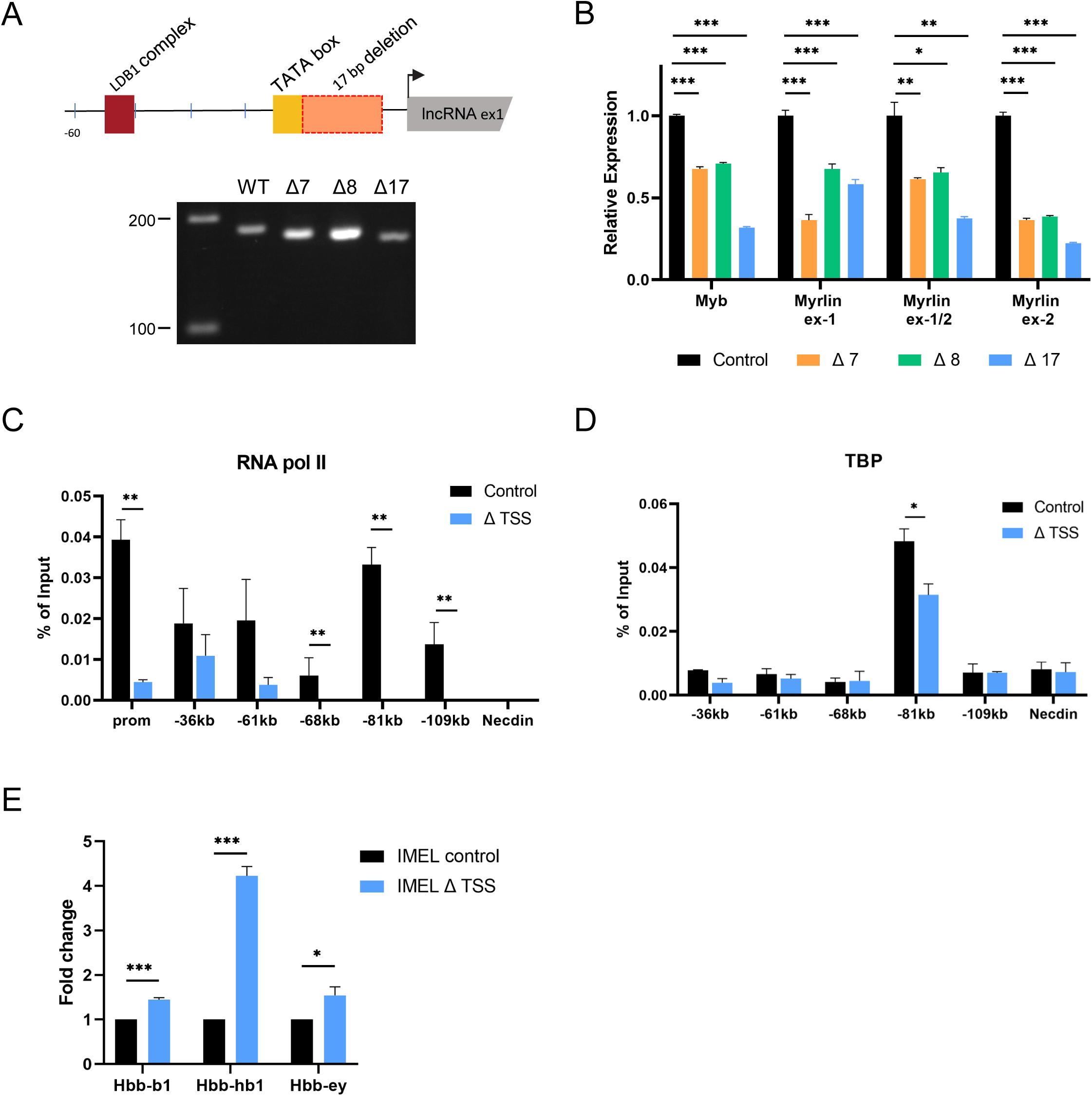
CRISPR deletions at the TSS for *Myrlin* affect *Myb* expression. (A) Schematic diagram of the CRISPR/Cas9-mediated 17bp deletion upstream of the *Myrlin* TSS. Smaller deletions were within the 17 bp deletion or extended upstream. PCR (below) shows the relative sizes for the 7bp, 8bp and 17bp deletion PCR products using WT or mutant gDNA. (B) Relative expression of *Myrlin* (exon 1, exon1/2 junction, exon 2) and *Myb* (exon 2) in MEL CRISPR/Cas9 control cells (no gRNA) and MEL 7bp, 8bp and 17bp CRISPR/Cas9 deletion mutants. Expression was normalized to *ActB*. (C) RNA Pol II-ChIP in MEL CRISPR/Cas9 control cells and MEL 17bp CRISPR/Cas9 deletion mutant (ΔTSS) at the *Myb* gene promoter and enhancer sites (−36, −61, −68, −71 and −109 kb). (D) ChIP for TBP at the *Myb* enhancer sites as in panel C for MEL CRISPR/Cas9 control cells and MEL ΔTSS CRISPR/Cas9 deletion mutant. (E) Relative expression of *Hbb-bh1*, *Hbb-b1* and *Hbb-y* in induced MEL CRISPR/Cas9 control cells and MEL ΔTSS CRISPR/Cas9 deletion mutant. Error bars indicate SEM of 3 independent biological experiments. (*) P < 0.05, (**) P<0.01, (***) P<0.001 by Student’s t-test. See also Figures S2.

To begin to characterize the impairment of *Myb* transcription ΔTSS cells, we performed ChIP to detect the occupancy of RNA Pol II. Compared to a control MEL cell line generated with a plasmid lacking an sgRNA, occupancy of Pol II in ΔTSS cells was reduced at the *Myb* promoter and at the −81 kb enhancer/*Myrlin* TSS as well as at the other enhancer sites (Figure 2C). TBP ChIP revealed that only the −81 kb enhancer was occupied by this member of the pre-initiation complex and in ΔTSS cells there was a marked reduction (Figure 2D). In accordance with results showing that fetal γ-globin transcription increases when *Myb* regulatory microRNAs are reduced in human cells ^27,44–46^, after differentiation of ΔTSS MEL cells, there was a several fold increase in murine embryonic βh1 globin transcription (Figure 2E), consistent with *Myb* reduction. These results support the idea that *Myrlin*, transcribed from the −81 kb *Myb* enhancer locus, is a positive regulator of *Myb* transcription.

### Long-range *Myb* promoter and enhancer contacts are reduced after *Myrlin* TSS deletion

*Myb* expression is regulated by long-range interactions between the *Myb* promoter and LDB1-bound enhancers within the *Myb-Hbs1l* intergenic region ^28^. A CTCF site 30 kb downstream of the *Myb* promoter also participates in the *Myb* enhancer hub, likely through direct interaction between CTCF and LDB1 ^47^. Moreover, the disruption of this enhancer hub underlies *Myb* downregulation during erythroid differentiation ^28^. We used chromatin conformation capture (3C) to determine whether ΔTSS influenced contacts between *Myb* and its intergenic enhancers. Compared to control cells, ΔTSS cells displayed reduced interaction frequency between the *Myb* promoter and enhancers, which was particularly evident at the −36 kb and −81 kb enhancer sites and the −30kb CTCF site (Figure 3A).

**Figure 3.**
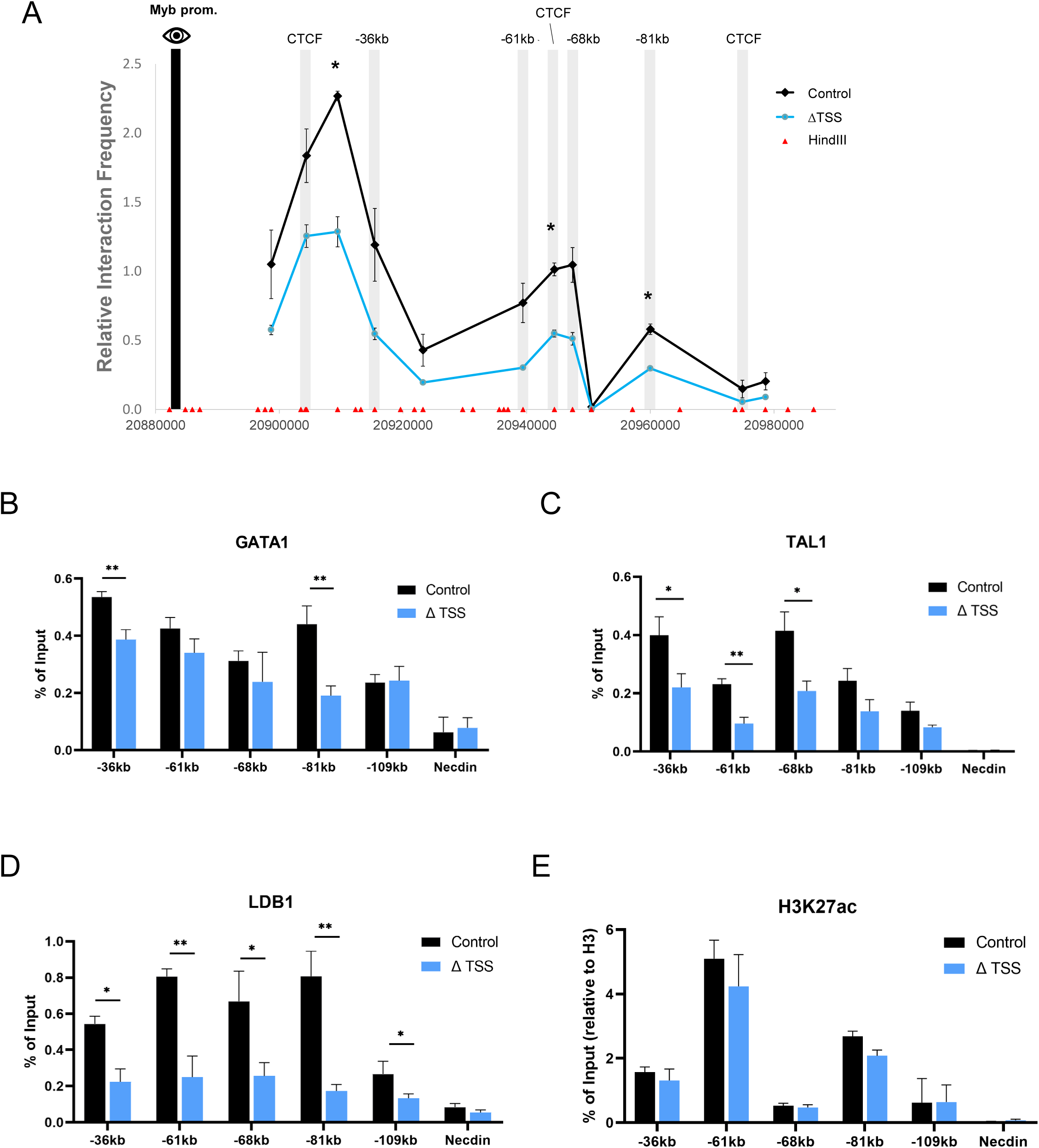
Chromatin organization of the *Myb* locus and transcription factor occupancy is affected by reduction of *Myrlin* in ΔTSS cells. (A) Chromatin conformation capture (3C) interaction frequency between the enhancers found within the *Myb*-*Hbs1l* intergenic region using the *Myb* promoter as the anchor (black bar) observed for ΔTSS and control cell uninduced MEL cells. (B) GATA1 occupancy at *Myb* enhancers in control and ΔTSS uninduced MEL cell lines determined by ChIP-qPCR. (C) TAL1 occupancy at *Myb* enhancers in control and ΔTSS uninduced MEL cell lines determined by ChIP-qPCR. (D) LDB1 occupancy at *Myb* enhancers in control and ΔTSS uninduced MEL cell lines determined by ChIP-qPCR. (E) H3K27ac normalized to H3 occupancy at *Myb* enhancers in control and ΔTSS uninduced MEL cell lines determined by ChIP-qPCR. Error bars indicate SEM of 3 independent biological experiments. (*) P < 0.05 and (**) P<0.01 by Student’s t-test.

The reduction of interaction frequency between *Myb* and its multiple enhancers by ΔTSS deletion closely resembles the reduced interactions observed upon LDB1 knock down using an shRNA in MEL cells ^28^. To investigate further, we carried out ChIP experiments to determine the occupancy of the LDB1 complex after reduction of *Myrlin* in ΔTSS cells. We observed that diminished long-range interactions in ΔTSS cells correlated with reduced LDB1 and TAL1 across the enhancers but reduced GATA1 occupancy was only notable at the −81 and −36 enhancers (Figure 3B-D). The H3K27ac mark, which indicates active enhancers, was not significantly affected in ΔTSS cells (Figure 3E). Together, our results show that the decrease in *Myb* expression upon deletion of the *Myrlin* TSS is accompanied by reduced LDB1 complex occupancy across the *Myb* enhancers and *Myb* enhancer hub disruption, although the enhancers remain in a potentially active state, retaining the H3K27ac mark.

### CRISPRi for *Myrlin* affects *Myb* transcription but not enhancer interactions

We next employed CRISPRi as an alternative approach to reduce the *Myrlin* transcript without altering the sequence context at the *Myb* −81 kb enhancer. Catalytically dead CAS9 (dCAS9) retains the ability to target DNA and can be an adaptable block to transcription elongation when fused to a KRAB repression domain. Using two different gRNAs to target dCAS9-KRAB to *Myrlin* exon 1, we observed a 50% to 60% reduction in *Myrlin* transcription leading to a similar drop in *Myb* transcription as observed in the *Myrlin* ΔTSS deletion (Figure 4A).

**Figure 4.**
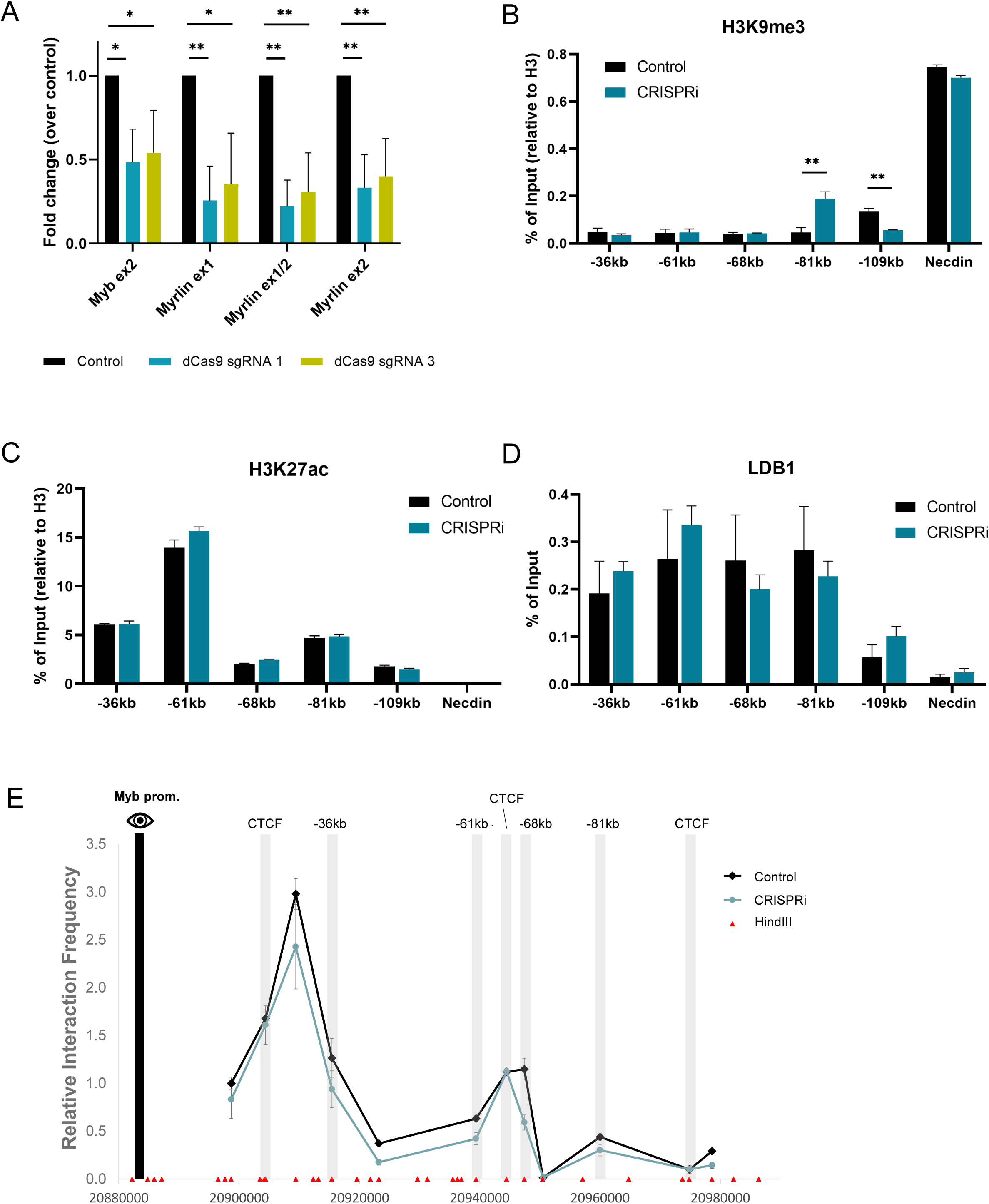
CRISPRi targeting of *Myrlin* compromises *Myb* transcription but not hub formation. (A) Expression of *Myb* and *Myrlin* monitored by RT-qPCR in CRISPRi uninduced MEL cells targeted with dCas9 sgRNA1, sgRNA3 or without an sgRNA (control). (B) ChIP-qPCR for H3K9me3 across the *Myb* locus before and after *Myrlin* CRISPi in uninduced MEL cells. (C) ChIP-qPCR for H3K27ac across the *Myb* locus before and after *Myrlin* CRISPi. (D) ChIP-qPCR for LDB1 across the *Myb* locus before and after *Myrlin* CRISPi. (E) Chromatin conformation capture (3C) interaction frequency between the *Myb* enhancers using the *Myb* promoter as the anchor (black bar) observed after *Myrlin* CRISPRi in uninduced MEL cells targeted with dCas9 sgRNA or without an sgRNA (control). Error bars indicate SEM of 3 independent biological experiments. (*) P < 0.05 and (**) P<0.01 by Student’s t-test.

ChIP localization of histone modifications at *Myb* enhancers revealed H3K9me3 at the −81kb enhancer after *Myrlin* CRISPRi, a signature heterochromatin mark of KRAB-mediated repression (Figure 4B). There was no change in H3K27ac at any enhancer after *Myrlin* CRISPRi, (Figure 4C), similar to what we observed after the *Myrlin* ΔTSS deletion. However, in contrast to broad reduction of LDB1 at enhancer sites after *Myrlin* ΔTSS deletion, LDB1 enhancer occupancy was not significantly reduced by *Myrlin* CRISPRi (Figure 4D). In addition, 3C experiments revealed relatively little change in interaction frequency between *Myb* and its enhancers after *Myrlin* CRISPRi compared to a control clone generated with a non-targeted dCAS9-KRAB vector (Figure 4E). Thus, downregulation of *Myb* after CRISPRi silencing of *Myrlin* does not involve loss of the *Myb* enhancer hub, separating looping and transcription activation at this locus. We conclude that *Myb* downregulation after *Myrlin* CRISPRi silencing does not require loss of the *Myb* enhancer hub, raising the possibility that *Myrlin* has a direct role in *Myb* transcription activation.

### *Myrlin* maintains H3K4me3 in the *Myb* locus through MLL1-WDR5

To further explore the role of *Myrlin* in *Myb* transcription activation, we focused on the *Myb* promoter and first exon/intron, which contain a CpG island that is highly enriched for H3K4me3 when *Myb* is active (Figure S3). ChIP-qPCR revealed strong H3K4me3 enrichment across these sequences, which was reduced following *Myrlin* transcriptional repression by *Myrlin* CRISPRi (Figure 5A). The Mixed Lineage Leukemia 1 (MLL1) complex has been shown to deposit H3K4me3 modifications at the *Myb* locus and activate transcription ^48^. Therefore, we carried out ChIP for MLL1 complex components MLL1 and WDR5. We found a strong decrease in both the MLL1 and WDR5 enrichment at the *Myb* locus after *Myrlin* CRISPRi (Figure 5B, C). Interestingly, HOTTIP long non-coding RNA targets the MLL1/WDR5 complex to promoters of HOXA genes to facilitate gene expression but currently no known eRNAs have been associated with MLL/WDR5 complex in the context of gene regulation ^23^.

**Figure 5.**
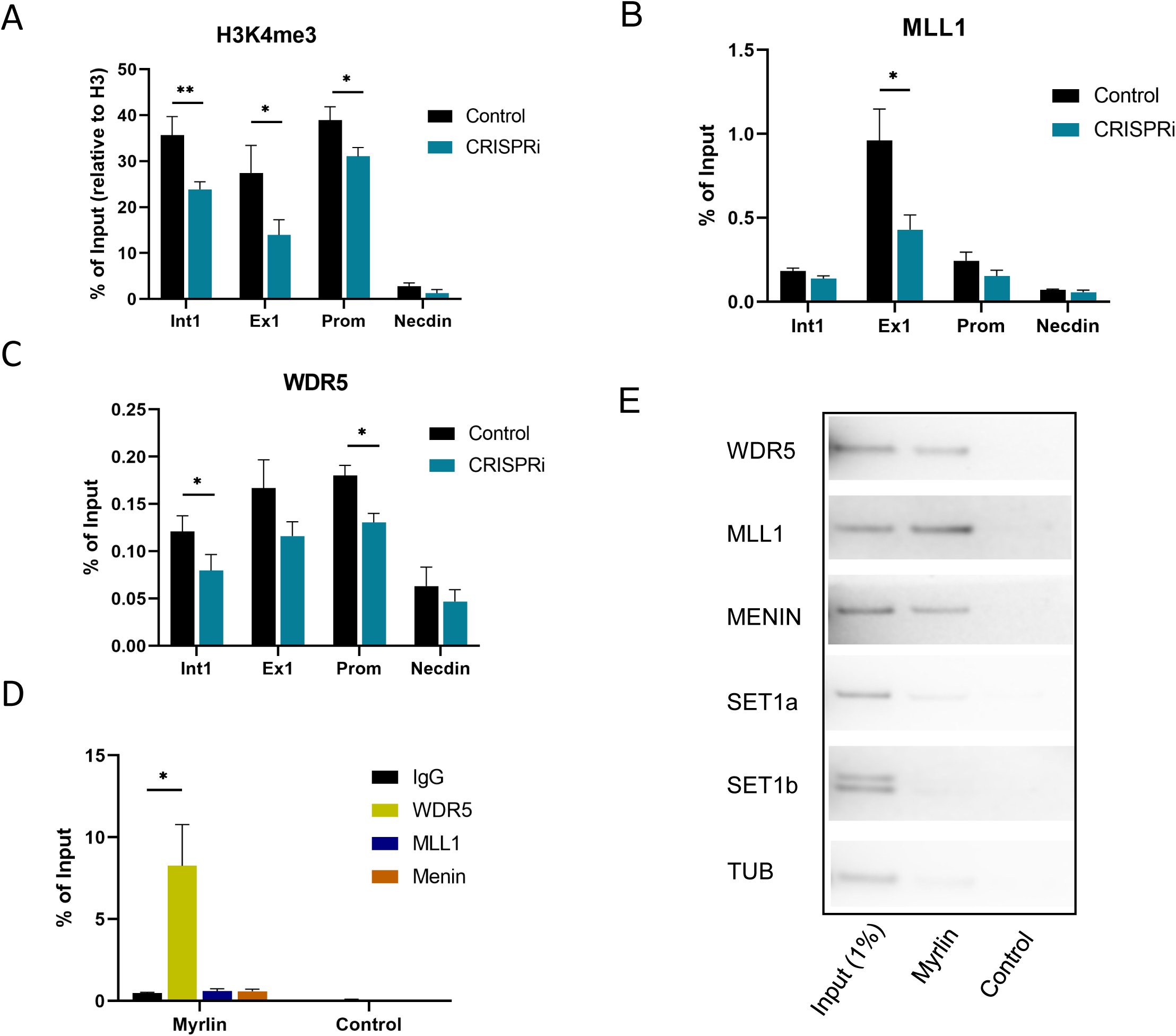
*Myrlin* interacts with MLL1 complex. (A) ChIP-qPCR for H3K4me3 across *Myb* sequences. (B) ChIP-qPCR for MLL1 at the *Myb* gene. (C) ChIP-qPCR for MLL1 complex component WDR5 at the *Myb* gene. (D) Myrlin RNA ChIP. (E) RNA pull down using biotinylated *Myrlin* and blotting using antibodies to MLL1 components. Tubulin served as a negative control. Error bars indicate SEM of 3 independent biological experiments. (*) P < 0.05 and (**) P<0.01 by Student’s t-test. See also Figures S3.

The results so far suggest that *Myrlin* may directly participate in activation of *Myb* transcription through recruiting MLL1. To further investigate, we performed *Myrlin* RNA-ChIP and found that the *Myrlin* transcript interacts with the MLL1 component WDR5 (Figure 5D). RNA-ChIP also detected interaction that were not statistically significant between *Myrlin* and MLL1 and with Menin, another component of the MLL1 complex. Therefore, we performed RNA pull down using biotinylated *Myrlin* and blotted against these components of the MLL1 complex. The results confirmed interaction of *Myrlin* with WDR5 and further revealed that *Myrlin* could pulldown complex components MLL1 and Menin. Biotinylated *Myrlin* did not pull down SET1a or SET1b, which are members of different COMPASS-like complexes, nor was there any interaction with Tubulin (TUB), which served as a negative control (Figure 5E). These results strongly support that *Myrlin* plays a role in MLL1 complex recruitment to promote H3K4me3 deposition and subsequent transcription of *Myb* and that *Myrlin* loss compromises this series of events and *Myb* transcription activation.

### RNA Pol II pausing and CDK9 occupancy are affected by CRISPRi of *Myrlin*

MLL1 complexes can maintain target gene expression through regulating both transcriptional initiation and elongation. MLL1 loss results in loss of the CDK9, the protein kinase subunit of pTEFb that confers phosphorylation on Pol II ser2 to drive transcription elongation in hematopoietic cells ^49,50^. Previous studies have shown that inhibition of CDK9 strongly reduced transcription elongation through the *Myb* gene body and resulted in pausing of Pol II within the *Myb* first intron ^28,51^. Using ChIP-qPCR, we found that CDK9 recruitment across the *Myb* enhancers and in *Myb* exon 1 and intron 1 was severely diminished after *Myrlin* CRISPRi (Figure 6A). Subsequent ChIP-qPCR analysis of RNA Pol II localization in the *Myb* locus revealed decreased occupancy at several *Myb* enhancers but no significant difference at the *Myb* promoter (Figure 6B). Interestingly, after *Myrlin* CRISPRi, RNA Pol II accumulated within the *Myb* gene body, particularly at *Myb* exon1/intron 1, which is consistent with reduction of *Myb* transcripts.

**Figure 6.**
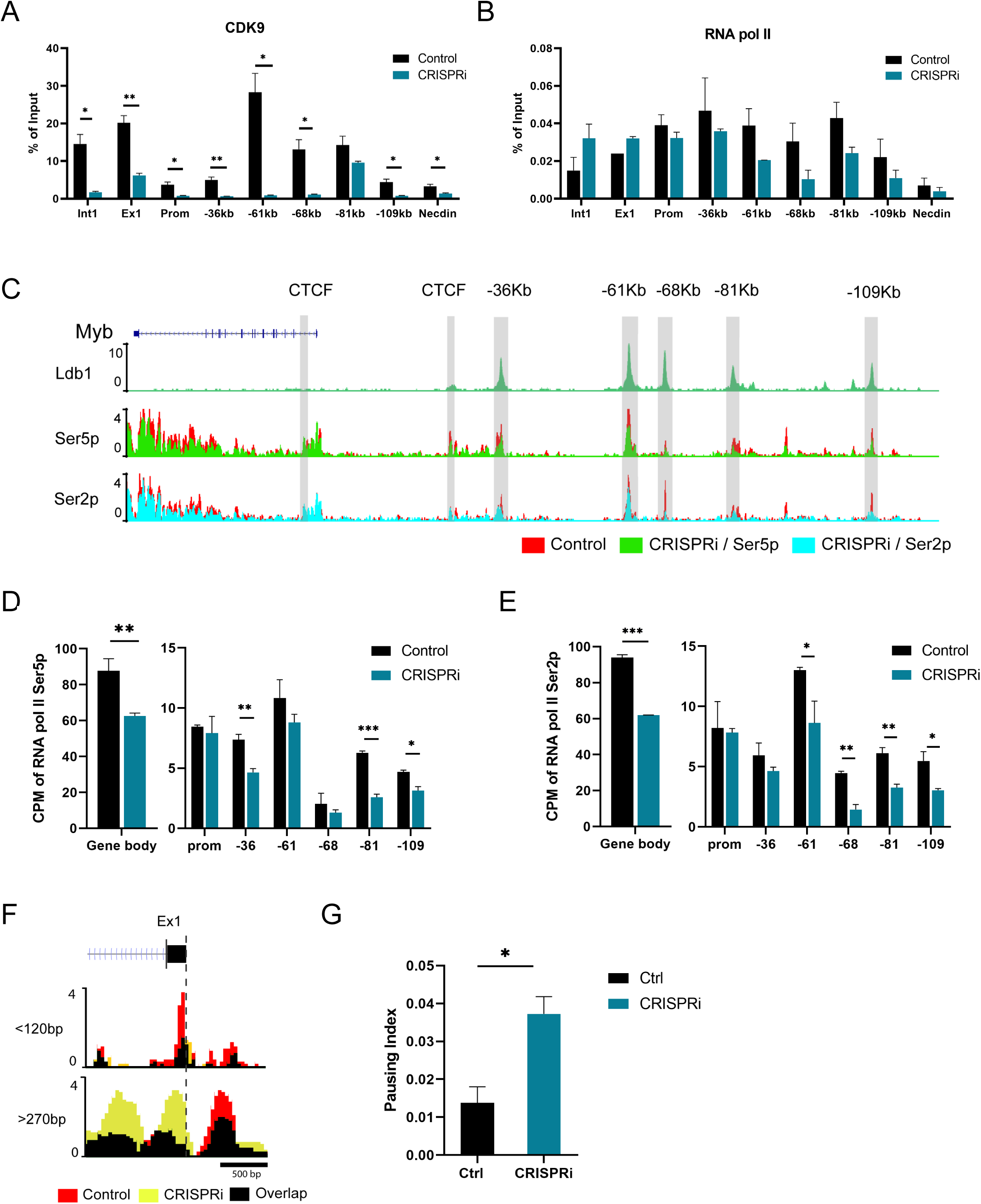
Pol II and CDK9 occupancy within the *Myb* enhancer hub is affected by CRISPRi targeting of *Myrlin*. (A) ChIP-qPCR for CDK9 in *Myrlin* CRISPRi uninduced MEL cells targeted with dCas9/KRAB or without an sgRNA (control). (B) RNA Pol II occupancy in the *Myb* promoter/exon 1 region after *Myrlin* CRISPRi and in control cells. (C) CUT&Tag for RNA Pol II Ser5 and Ser2 phosphorylated forms. (D, E) Quantitation of CUT&TAG data showing Pol II Ser5P and Ser2P occupancy in the *Myb* gene body and at each of the enhancers. (F) Separate analysis of the shorter (<120 bp) and longer (>270 bp) Pol II Ser5P CUT & TAG fragments displaying Pol II Ser5 occupancy ^52^. (G) Pausing index calculated for Pol II Ser5 across *Myb*. Error bars indicate SEM of 2 independent biological experiments. (*) P < 0.05 and (**) P<0.01 by Student’s t-test. See also Figures S4.

To further investigate this result, we performed CUT & TAG for the Ser5P and Ser2P phosphorylated forms of RNA Pol II, which represent the initiating and elongating forms of the enzyme, respectively. The genome browser view in Figure 6C illustrates the reduction of both phosphorylated forms of RNA Pol II across the *Myb* locus after *Myrlin* CRISPRi. Quantitation of these data shows about a 30% reduction of Pol II Ser5P and Ser2P within the *Myb* gene body and reduced occupancy at each of the enhancers (Figure 6D, E). To obtain increased resolution of this result, we analyzed the signals of the shorter (<120 bp) and longer (>270 bp) Pol II Ser5P CUT & TAG fragments, which distinguishes Pol II in the pre-initiation state at the TSS and paused Pol II at peaks up- and downstream of the TSS, respectively ^52^ (Figure 6F). *Myrlin* CRISPRi reduced the signal for the shorter fragments, suggesting some decrease in Pol II recruitment upon *Myrlin* loss. At the same time, the signal associated with longer fragments, indicative of paused Pol II, strongly increased across exon/intron 1.

We calculated the pausing index of Pol II Ser5P from the longer fragment data based on the Ser5P ratio at the TSS (exon1) and in the gene body and found about a 3-fold higher pausing index in *Myb* after *Myrlin* CRISPRi compared to controls (Figure 6G). Similar results were obtained for Pol II pausing in *Myb* using RNAseq peaks without separation by fragment size (Figure S4A, B). These results connect *Myrlin* to the mechanism by which transcription of *Myb* is regulated and suggest that *Myrlin*, through MLL1, may help to recruit or stabilize CDK9 at *Myb,* which is necessary for efficient RNA Pol II elongation through the gene.

### KLF1 interacts with *Myrlin* transcripts within the *Myb* active chromatin hub

What could be the basis of *Myrlin* local function in *Myb* transcription? *Myrlin* interaction with MLL1 at the actively transcribed *Myb* promoter raised the question of whether *Myrlin* is retained within the enhancer hub by tethering to the −81 kb enhancer, adjacent to its transcription start site. We performed ChIRP experiments to determine *Myrlin* localization in the *Myb* locus. Two sets of *Myrlin* probes (odd and even probes) can specifically capture the *Myrlin* transcript (Figure 7A). We found prominent peaks for *Myrlin* at its two exons and at the −81 kb enhancer site but not at other sites across the locus (Figure 7B). Since the −81 kb enhancer is close to *Myrlin* exon 1, the signal at that site could have been contributed to by cross-linking of *Myrlin* bound to exon1. Therefore, we asked whether *Myrlin* interacted with −81 kb-bound transcription factors, specifically KLF1, which uniquely binds to the −81 site among the enhancers. KLF1 is required for full *Myb* transcription activation ^28^. Moreover, transcription factors have recently been documented to commonly bind RNAs ^53^. Indeed, RNA ChIP for KLF1 revealed interaction with *Myrlin*, which was confirmed by biotinylated RNA pull down (Figure 7C, D). ChIP-qPCR confirmed occupancy of KLF1 at the −81 kb *Myb* enhancer but not at −68 kb, as expected (Figure 7E). KLF1 occupancy was not affected by *Myrlin* CRISPRi. We conclude that KLF1 may contribute to the localization of *Myrlin* within the *Myb* enhancer hub.

**Figure 7.**
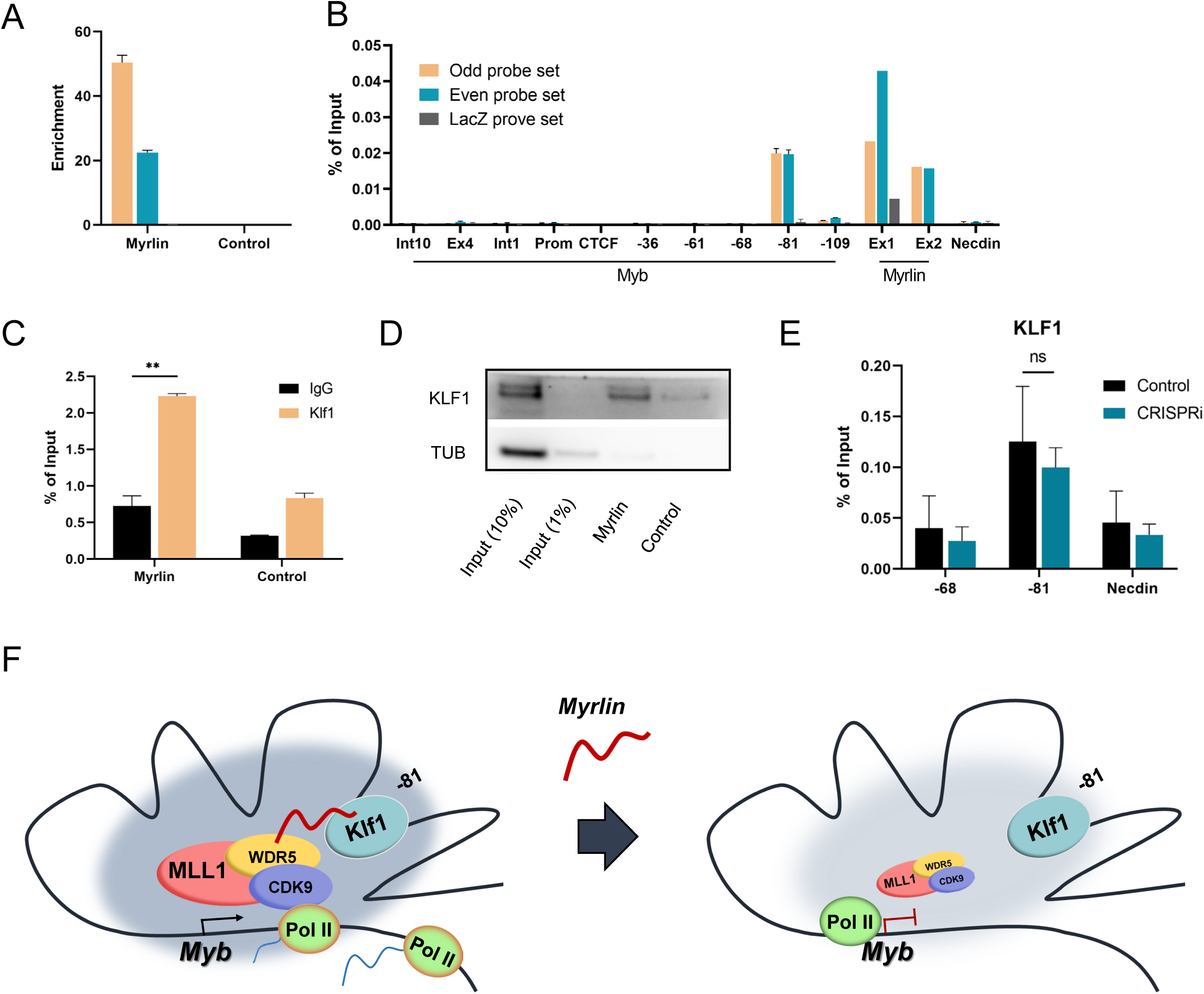
KLF1 interaction with *Myrlin* contributes to localization within the *Myb* enhancer hub. (A) *Myrlin* RNA pull down was conducted to determine efficiency of probes for ChIRP. (B) ChIRP DNA pull down by *Myrlin* across *Myb* and the *Myb* enhancers. (C) RNA ChIP for KLF1. (D) Biotinylated *Myrlin* pull-down and blotting with KLF1 antibodies. (F) Model of the *Myb* locus enhancer hub. Transcription of *Myb* is depicted with and without the −81 kb enhancer *Myrlin* eRNA after *Myrlin* CRISPRi. Large, shaded circle represents Pol II and LDB1 transcription factor density within the *Myb* enhancer hub which is diminished when *Myrlin* transcription is reduced. Error bars indicate SEM of 3 independent biological experiments. (*) P < 0.05 and (**) P<0.01 by Student’s t-test.

In summary, structural components contributing to *Myb-*enhancer interactions within the hub, including the LDB1 complex and KLF1, are present and the hub forms normally when *Myrlin* is lost through CRISPRi (Figure 7F). However, in the absence of *Myrlin*, recruitment of MLL1/WDR5 is reduced, resulting in poor binding of CDK9 and RNA Pol II in the *Myb* locus. Moreover, RNA Pol II pausing within the *Myb* first exon/intron and failure to complete *Myb* transcripts are observed upon *Myrlin* loss. We conclude that the *Myrlin* eRNA has important recruitment functions for MLL1 complex that are required for *Myb* transcription activation beyond formation of the *Myb* enhancer hub.

## DISCUSSION

Hundreds of long non-coding RNAs are expressed during erythropoiesis but evidence for their function remains anecdotal ^33–35,37^. We identified *Myrlin*, an eRNA, transcribed from the −81 kb murine *Myb* enhancer as a positive regulator of *Myb*. A rigorous paradigm to deduce the function of lncRNA and eRNA loci involves dissection of the functional roles of the RNA transcript versus the act of enhancer transcription, both of which can influence target gene expression and/or chromatin organization ^54^. To this end, we analyzed the role of the RNA molecule itself by using *Myrlin* RNA ChIP and biotinylated *Myrlin* pull down assays to document the importance of the transcript, *per se*. We found that *Myrlin* interacts with MLL1/WDR5 to promote *Myb* transcription. In the absence of *Myrlin*, MLL1 recruitment to *Myb* and *Myb* promoter H3K4me3 modification is reduced. In addition, CDK9 is reduced at the *Myb* promoter and RNA Pol II accumulates excessively within the *Myb* first exon/intron. Thus, *Myrlin* is required for Pol II pause release to promote *Myb* transcription.

In our initial experiments to reduce *Myrlin* transcription, we deleted 17 bp of the *Myrlin* TSS. This resulted in loss of the LDB1 looping complex from all the *Myb* enhancer sites, disruption of the *Myb* enhancer hub and reduction of *Myb* transcription. The results suggest interdependence among multiple *Myb* enhancers for formation of the hub. Hubs are understood from single allele interaction experiments to represent close association of all hub sequences within which multiple enhancers show preferential aggregation with each other and with the genes they regulate ^55^. Enhancer interdependence has been observed at the *IgK* super enhancer cluster where deletion of one enhancer affects interactions among the others ^56,57^. This model comports well with the idea that extrusion of chromatin loops through the action of cohesin complexes and CTCF will bring many points within an extended locus into close proximity where their interactions may be stabilized by specific transcription factors ^58,59^. Such a hub of interactions and transcription factor density may facilitate accumulation of transcriptional components such as RNA Pol II, possibly involving liquid-liquid phase separation, favoring transcription ^14^.

The precise role of *Myrlin* in *Myb* enhancer hub formation, if any, remains to be clarified, since the LDB1 complex was lost from the enhancer sites in the ΔTSS cells, which is well documented to compromise LDB1 enhancer looping in this locus ^28^. Intriguingly, LDB1 binding, and the hub formation were not affected when *Myrlin* transcription was repressed by CAS9-KRAB, which did not affect LDB1 complex occupancy. It remains unclear why LDB1 is lost from the enhancer sites in ΔTSS cells but not after *Myrlin* CRISPRi. One possibility is that the 17 bp deletion of the TSS, near the LDB1 complex motif at the −81 kb enhancer, distorts the local chromatin architecture sufficiently to reduce LDB1 binding, which then destabilizes binding at the other hub enhancers, and looping. Regardless, losing *Myrlin* under the circumstances of KRAB repression did not have this effect on LDB1 binding yet still resulted in *Myb* downregulation, separating looping from transcription activation. Looping in the absence of transcription has been observed at the β-globin locus in erythroid cells and, more generally, in leukemia cells after drug treatment to inhibit BET proteins ^60–62^. These data support the idea that enhancer looping mechanistically precedes transcription activation.

To explore the difference between looped and transcriptionally active *Myb* loci versus inactive looped loci, we localized RNA Pol II and found a decrease in Pol II recruitment to the *Myb* promoter after *Myrlin* CRISPRi and significant pausing in intron/exon 1. Pol II pausing is correlated with transcriptional repression during terminal erythroid repression ^63^. Previous work had suggested a Pol II pause region in *Myb* intron 1 that was proposed to correspond to either a stem-loop forming sequence followed by a poly (dT) tract 1.7 Kb downstream of the *Myb* TSS or to a CTCF site about 2 Kb downstream ^28,51^. These groups reported that transcription elongation through *Myb* was inhibited in erythroid cells or in breast cancer cells by CDK9 inhibitors. Our high-resolution CUT & TAG results for Pol II Ser5 and Pol II Ser3 showed localization of paused transcripts across *Myb* exon 1 and into intron 1, likely encompassing the sites previously suggested as pause sites. We were able to show that *Myrlin* functions to recruit or stabilize CDK9 in the *Myb* exon/intron1 Pol II pause region. It has been suggested that interactions of an enhancer with its target promoter can stimulate Pol II pause release (Ghavi-helm 2014, Rahl 2010) and that this property may be related to CDK9 activity ^64–66^. *Myrlin* provides a link between an eRNA and CDK9 at a target gene.

We observed that *Myrlin* interacts with several subunits of the MLL1 complex, including MLL1, WDR5 and Menin, and that *Myrlin* is important for MLL1 occupancy in the *Myb* locus and for H3K4me3 modification at the *Myb* promoter CpG island. Promoter proximal H3K4me3 has recently been linked to RNA Pol II pause release ^67^. Reduced MLL1 may underlie loss of CDK9 at the *Myb* locus, as CDK9 reduction was reported on a global scale after deletion of MLL1 in hematopoietic stem and progenitor cells ^49^. In this scenario, the *Myrlin* eRNA becomes important for *Myb* transcription after the enhancer hub is formed. We propose that *Myrlin* interacts with MLL1 and the CDK9 component of pTEFb to recruit RNA Pol II to the locus and assure efficient elongation through the *Myb* exon/intron 1 pause region. After *Myrlin* CRISPR, MLL1 and CDK9 are reduced, and Pol II pausing reduces *Myb* transcripts. The above putative functions of *Myrlin* would require its localized presence in the *Myb* locus/enhancer hub. We explored the potential role of KLF1 in such localization since it binds uniquely to the −81 enhancer where the *Myrlin* eRNA is transcribed. Indeed, we documented interaction of *Myrlin* and KLF1 by biotinylated RNA pull down.

One of the most promising strategies for treating Sickle Cell Disease and β-Thalassemia is reactivation of fetal hemoglobin production in erythroid cells of adult patients. GWAS revealed an association between single nucleotide polymorphisms (SNPs) in the *Myb* ─ *Hbs1l* intergenic region, encompassing the several *Myb* enhancers, that reduce *Myb* gene expression and elevate fetal hemoglobin in human adult erythroid cells ^26,29,30^. Two of these SNPs, i.e., rs66650371 and rs77755698, are located within the LDB1 complex GATA1/TAL1 binding peak at the human −84 kb *Myb* enhancer, which is the homologue of the murine −81 kb enhancer ^29,68^. These SNPs reduce LDB1 binding to the −84 kb enhancer and decrease interaction frequency with the *MYB* promoter ^29^. The effect of the SNPs on interactions of the other enhancers with MYB was not tested in this work but our results suggest that overall formation of the hub is likely affected.

Recently, a 1,283 bp non-coding RNA, HMI-LNCRNA, was reported to arise from the −84 kb human *MYB* enhancer ^36^. Thousands of human lncRNAs have homologues in other species with similar expression patterns but low sequence conservation ^35,69^. However, BLATN sequence comparison of HMI-LNCRNA to the mouse genome revealed a conserved ‘patch’ of 378 nt (85.8% homology) shared with the 5’ end of *Myrlin*, something commonly observed for these poorly conserved lncRNAs ^69^ (Figure S5). Within this homology patch lie the GATA1/LDB1 complex binding site that mediates looping to the *Myb* promoter in human and mouse erythroid cells, the −81 kb KLF1 binding site and the *Myrlin* TATA box and first exon. Thus, it seems likely that the pause release function of *Myrlin* at the *Myb* gene may be conserved between species. While any *Myb* regulatory function ascribed to *Myrlin* in mouse cells remains to be established for the related transcript in human cells, the *Myb* transcription mechanisms participated in by *Myrlin* suggest the possibility of their utility as targets to increase HbF production to ameliorate the severity of hemoglobinopathies.

## AUTHOR CONTRIBUTION

AD and IK conceived the project; IK, JK and BL supervised experiments. LFD, JK and MJM performed experiments. LFD, JK and AD wrote the paper; all authors edited the paper.

## ACKNOWLEDGEMENTS

We acknowledge computational support from Dr. Lecong Zhou, the NIDDK Genomics Core for sequencing support and the NIH HPC (Biowulf) for computational support. This work was funded by the Intramural Program of the National Institute of Diabetes and Digestive and Kidney Diseases, NIH (DK 075033 to AD). We thank Dr. Xiang Guo for kindly providing the mouse tissues.

## DECLARATION OF INTERESTS

The authors declare that they have no conflicts of interest.

## RESOURCE AVAILABILITY

### Lead contact

Further information and request for resources and reagents should be directed to and will be fulfilled by the Lead Contact, Ann Dean.

### Materials availability

Plasmids generated in this study are available by request from the lead contact.

### Data and code availability

- Sequencing data discussed in this publication are deposited in the Gene Expression Omnibus with accession code GSE240060.
- ENCODE data can be obtained from integrative publication PMID: 22955616; PMC: PMC3439153.
- This paper does not report original code.
- Any additional information required to reanalyze the data reported in this paper is available from the lead contact upon request.

## EXPERIMENTAL MODEL AND STUDY PARTICIPANT DETAILS

### Cell culture and animals

Control and CRISPR-edited mouse erythroid leukemia (MEL) cell lines were cultured in a 5% CO2 humidified incubator at 37°C in DMEM (Gibco, 11965118) with L-Glutamine (Gibco, A2916801), 10% fetal bovine serum (R&D system, S12450), 1% Penicillin and Streptomycin (Gibco, 15070063) and 1mM sodium pyruvate (Gibco, 11360070) at a density between 1×10^5^ and 1×10^6^ cells/mL. MEL differentiation was induced at a concentration of 2.5 x 10^5^ cells/mL with 2% DMSO (Millipore sigma, D4540) for 4 days.

### Mice and ethics statement

Mouse protocols were approved by the NIDDK Animal Care and Use Committee in accordance with AALAC specifications. Yolk sacs from E8.5 and fetal livers from E14.5 were dissected and washed in phosphate-buffered saline.

## METHOD DETAILS

### CRISPR-Cas9 genome editing of MEL cells

CRISPR gRNAs were designed using GeneTargrter (http://genetargeter.mit.edu/) (see Table S1 for gRNA sequences). gRNAs targeting the *Myrlin* transcription start site were cloned into the CRISPR-Cas9 and gRNA expression vector pSpCas9(BB)-2A-GFP (PX458) (gift from Feng Zhang, Addgene plasmid #48138) as described ^70^. MEL cells were transfected with Escort IV lipid transfection reagent (Sigma-Aldrich L3287) according to the manufacturer’s instructions. Fluorescent cells were sorted 48 hours later and plated at limiting dilution to isolate clones. Clonal lines were genotyped by PCR using EmeraldAmp GT PCR Master Mix (Takara, RR310A) and target specific primers flanking the *Myrlin* transcription start site. Deletions were validated by sequencing.

Stable MEL cell clones expressing dCAS9-KRAB were generated using Lenti-dCAS9-KRAB-blast (Gift from Dr. Gary Hon, Addgene plasmid #89567). 2 × 10^6^ of MEL cells were suspended in 100ul of High-Performance Electroporation Solution (BTXpress, 45-0801) with 5-10 µg of plasmid DNA and electroporated with the Gene Pulser Xcell System (Bio-Rad) using 2 pulses at 200 V for 5 ms. Cells were diluted with 100 µL of pre-warmed media and transferred to 2 mL of media in a 12-well culture dish. Cells were selected in 10 µg/mL Blasticydin (Gibco, A1113903) for one week and plated at limiting dilution to isolate clones. Clonal lines were checked for production of S. pyogenes dCAS9 by RT-qPCR. dCAS9-KRAB MEL cells were electroporated as above with gRNAs targeting the *Myrlin* transcription start site cloned into LentiGuide-puro (gift from Feng Zhang, Addgene plasmid #52963). Cells were selected in 10 µg/mL Blasticydin and 1 µg/mL puromycin (Gibco, A1113803) for one week and plated at limiting dilution to isolate clones. Transfected cells were validated for expression of *Myrlin* by RT-qPCR.

### 5’ and 3’ rapid amplification of cDNA ends (RACE)

RACE was performed using the FirstChoice RLM-RACE kit (ThermoFisher Scientific, AM1700) following the manufacture’s protocol. Total RNA from MEL cells was extracted and reverse transcribed using the 3’ RACE adapter and the sequence of interest was amplified by nested PCR (3’ RACE). Alternatively, a sample of the same RNA was treated with Calf Intestine Alkaline Phosphatase, then Tobacco Acid Pyrophosphatase and finally ligated to the 5’ RACE adapter. De-capped adapter-ligated RNA was then reverse transcribed, and the sequence of interest was amplified by nested PCR (5’ RACE). PCR products were separated on a 1% agarose gel, purified using the QIAquick gel extraction kit (Qiagen, 28704), and cloned into the pCR4-TOPO vector (Invitrogen, K457502) for sequencing. For RACE primers, see Table S1.

### Reverse-transcription qPCR (RT-qPCR)

RNA was isolated from 1×10^6^ MEL cells with the RNeasy Plus kit (Qiagen, 74134). RNA was reverse-transcribed using the Superscript III First-Strand Synthesis System (ThermoFisher Scientific, 18080051) following manufacturer’s instructions. RT-qPCR was performed using the iTaq Universal SYBR Green Supermix (Bio-Rad, 1725120) with the ABI 7900HT (Applied Biosystems). Data was normalized to ActB. For RT-qPCR primers see Table S1 and Stadhouders et al., 2012 ^28^.

### Chromatin immunoprecipitation (ChIP)

10^6^ of MEL cells per IP were cross-linked with 1% formaldehyde in PBS at room temperature for 10 minutes. Glycine was added to a final concentration of 0.125 M and sample were incubated at room temperature for 5 minutes. Cells were washed three times with cold PBS by centrifugation. For LDB1 ChIPmentation cells were double cross-linked. MEL cells were washed three times with cold PBS with 1 mM MgCl_2_. Disuccinimidyl glutarate (ThermoFisher Scientific, 20593) in DMSO were added to a final concentration of 2 mM and samples were rocked at room temperature for 45 minutes. Cells were washed three times with PBS by centrifugation and cross-linked with 1% formaldehyde in PBS at room temperature for 10 minutes, Glycine weas added to a final concentration of 0.125M and sample were incubated at room temperature for 5 minutes. Cells were washed three times with cold PBS by centrifugation. Cell pellets were snap frozen and thawed on ice on the day of starting the ChIPmentation experiment. Cell pellets were resuspended in 1 mL ChIP sonication buffer (10 mM Tris pH 8.0, 0.25% SDS, 2 mM EDTA) with proteinase inhibitors (Sigma, P8340) and 1 µM phenylmethylsulfonyl fluoride (PMSF) and sonicated by 12 cycles (30% amplitude, 30 sec on/30 sec off) using Sonifier SFX250 (BRANSON). Then samples were diluted in 1:1.5 ratio with equilibration buffer (10 mM Tris, 233 mM NaCl, 1.66% TritonX-100, 0.166% Na-Deoxycholate, 1 mM EDTA, proteinase inhibitors and 1 µM PMSF). Samples were spun at 14,000g for 10 minutes at 4°C and supernatant was transferred to a new tube. 1% input was preserved. 3 µg of antibody of interest or isotope-matched IgG was added, and samples were incubated on a rotator overnight at 4°C. On the second day, Pierce™ ChIP-grade Protein A/G Magnetic Beads (ThermoFisher Scientific, 26162) were washed once with RIPA-LS (10 mM Tris pH 8.0, 140 mM NaCl, 1 mM pH 8.0 EDTA, 0.1% SDS, 0.1% Na-Deoxycholate, 1% TritonX-100, proteinase inhibitors and 1 µM PMSF) and added to chromatin. ChIP reactions by adding 25 µL /IP of A/G beads rotated for 2 hours at 4°C. Chromatin bound beads were washed twice with ice cold RIPA-LS, twice with ice cold RIPA-HS (10 mM Tris pH 8.0, 500 mM NaCl, 1 mM pH 8.0 EDTA, 0.1% SDS, 0.1% Na-Deoxycholate, 1% TritonX-100, proteinase inhibitors and 1 µM PMSF), twice with ice cold RIPA-LiCl (10 mM Tris pH 8.0, 250 mM LiCl, 1 mM pH 8.0 EDTA, 0.5% NP-40, 0.5% Na-Deoxycholate, proteinase inhibitors and 1 µM PMSF) and once with TE (10 mM Tris pH 80, 1 mM EDTA pH 8.0). Beads were resuspended with 48 µL of ChIP elution buffer (10 mM Tris pH 8.0, 300 mM LiCl, 5 mM pH 8.0 EDTA, 0.4% SDS) and 2 µL of proteinase K (20mg/mL, Invitrogen, AM2546). 0.4% SDS, 300 mM NaCl and 2 µL of proteinase K were added for input samples. Samples were incubated at 55°C for one hour, then 65°C for 6-10 hours. DNA was purified using the ChIP DNA Clean & Concentrator Kit (Zymo Research, D5205) according to the manufacturer’s instructions. Real-time qPCR was performed with published primers ^28^ using the iTaq Universal SYBR Green Supermix. Percent of input was calculated against input. For ChIPqPCR primers and antibodies, see Table S1.

### Chromatin conformation capture assay (3C)

Cells were cross-linked with 2% formaldehyde in PBS at room temperature for 5 minutes, Glycine weas added to a final concentration of 0.125 M and sample were incubated at room temperature for 5 minutes. Cells were washed twice with cold PBS and resuspended in lysis buffer (10 mM Tris-HCl pH 8.0, 10 mM NaCl, 0.2% NP40, proteinase inhibitor). After lysis for 30 minutes on ice, nuclei were collected by centrifugation at 800g for 10 minutes at 4°C and resuspended in 1.2 X of NEB buffer 2.1 (New England Biolabs, B7202). 1 x 10^7^ nuclei were then solubilized with 0.3% SDS for one hour at 37°C followed by adding 1.8% of TritonX-100 and incubating one hour at 37°C. Chromatin was digested with 1000 U of HindIII (New England Biolabs, R0104M) overnight at 37°C. Digested genomic DNA control were taken and stored at −20°C. Restriction enzyme was inactivated using 1.6% of SDS for 25 minutes at 65°C and chromatin was ligated by adding 4000 U of T4 ligase (New England Biolabs, M0202M) in 1× T4 DNA ligase buffer (New England Biolabs, B0202S) containing 1% TritonX-100 and for 4 hours at 16°C followed by additional incubation for 30 minutes at room temperature. Chromatin was de-cross-linked with a final concentration of 100 µg/mL of proteinase K at 65°C for 6-10 hours. Then 0.5 µg/mL of RNase A was treated for one hour at 37°C. Samples were purified via phenol-chloroform extraction and ethanol precipitation. Relative cross-linking between the *Myb* promoter and fragments of interest was analyzed by real-time qPCR with published TaqMan probes and primers ^28^. Ligation products of HindIII digested BAC DNA containing the mouse *Myb*-*Hsbl1* intergenic region were used to determine primer efficiency. Additional primers are listed in Table S1.

### *Myrlin* RNA cellular localization

MEL cells were washed with ice cold PBS and lysed in hypotonic buffer (25 mM HEPES, 2 mM EDTA, 0.5% Tween-20). Cytoplasmic and nuclear fractions were obtained by centrifugation at 800 g for 10 minutes at 4°C. RNA from the supernatant cytoplasmic material and the nuclear pellet were purified and RNA was reverse transcribed, and cDNA was measured by real-time qPCR. For RT-qPCR primers see Table S1.

### Chromatin Isolation by RNA purification (ChIRP)

The *Myrlin* probes were designed using the Stellaris Probe Designer version 4.2 and synthesized with 3’ Bio-TEG modification by IDT. LacZ probes were used as a negative control (Millipore, CS216572). ChIRP-qPCR was performed as described (Chu *et al.*, 2012). 2 × 10^6^ of MEL cells per IP were fixed with 1% glutaraldehyde (Sigma-Aldrich, G5882) for 10 minutes at room temperature. Glycine was added to a final concentration of 0.125 M and sample were incubated at room temperature for 5 minutes. Cells were washed three times with cold PBS by centrifugation. Cell pellets were snap frozen and thawed on ice on the day of starting the ChIRP experiment. Cell pellets were resuspended in lysis buffer (10 X the mass of pellet, 50 mM Tris pH 7.0, 1% SDS, 10 mM EDTA) with proteinase inhibitors, 0.1 U/µL Superase-in (Invitrogen, AM2694) and 100 mM PMSF, then subjected to sonication with Bioruptor (Diagenode) for 50 minutes at highest setting with 30 seconds ON, 45 seconds OFF pulse intervals. Samples were spun at 16,000 g for 10 minutes at 4°C and supernatant was transferred to a new tube. Chromatin was snap frozen and thawed on ice on the day of starting the ChIRP experiment. Chromatin is diluted in two times volume of hybridization buffer (750 mM NaCl, 1% SDS, 50 mM Tris pH 7.0, 1 mM EDTA, 15% Formamide) with proteinase inhibitors, 0.1 U/µL Superase-in and 100 mM PMSF. Pre-designed probes were separated into two groups which are even probe set (probe 2, 4, 6 and 8) and odd probe set (probe 1, 3, 5 and 7) and each probe set (100 pmol /1 mL chromatin) were added to diluted chromatin, which was mixed by end-to-end rotation for 4 hours at 37°C. Dynabeads MyOne Streptavidin C1 (Invitrogen, 65001) were washed three times in lysis buffer and resuspended in its original volume. 100 µL of streptavidin beads were added per 100 pmol of probes, and samples were mixed for 30 minutes at 37°C. Chromatin bound beads were washed five times with 1 mL of pre-warmed wash buffer (2× SSC, 0.5% SDS, 100 mM PMSF) for 5 minutes at 37°C. At last wash, 100 µL of resuspended beads aliquoted for RNA isolation and 900 µL for DNA fraction. RNA was extracted with the miRNAeasy Mini Kit (Qiagen, 217004), and Superscript III First-Strand Synthesis System was used to reverse transcribe RNA to cDNA. DNA fraction was isolated by phenol-chloroform extraction and ethanol precipitation. qPCR was performed using iTaq Universal SYBR Green Supermix with the ABI 7900HT.Percent of input was calculated against input. For probes see Table S1.

### RNA pulldown

Full-length *Myrlin* RNA was generated using the MEGAscript T7 Kit (Invitrogen, AM1333) according to the manufacturer’s protocol. RNA pulldown was performed using the Pierce Magnetic RNA-Protein Pull-Down Kit (ThermoFisher Scientific, 20164) according to the manufacturer’s protocol. Briefly, 1 μg of biotinylated *Myrlin* RNA was incubated with 1 mg of precleared protein extracted from MEL cells for 4 hours at 4°C. Following this, Streptavidin magnetic beads was added and incubated for an additional 2 hours. Finally, proteins were eluted and subjected to western blotting.

### Western blot

Proteins were separated on NuPAGE 4%–12% Bis-Tris Gel (Thermo Fisher Scientific, NP0321) and transferred to a PVDF membrane using Trans-Blot Turbo Transfer Pack (Bio-RAD, 1704156). Membrane was blocked in blocking buffer (1 X TBS, 0.1% Tween-20, 5% w/v Nonfat dry milk) for 1 hour at room temperature. Primary antibodies in blocking buffer were treated overnight at 4°C. Membranes were washed with TBST three times with shaking for 10 minutes, incubated with HRP-conjugated secondary antibody in blocking buffer for 1 hour at room temperature, and washed three times in TBST. Blots were exposed to SuperSignal West Dura Extended Duration Substrate (Thermo Fisher Scientific, 34075) and scanned by Syngene PXi (Syngene). Antibodies are listed in Table S1.

### RNA-ChIP

The RNA ChIP-IT Kit (Active Motif, 53024) was used according to the proprietary methods. 10^6^ of MEL cells were collected and cross-linked with 1% formaldehyde at room temperature for 10 minutes. For KLF1 RNA-ChIP cross-linking was not applied since KLF1 binding site at −81 kb *Myb* enhancer is close to *Myrlin* gene location. Nuclei were isolated and sonicated by 10 cycles (30% amplitude, 30 sec on/30 sec off) using Sonifier® SFX250. Immunoprecipitated RNA using antibody of interest or isotope-matched IgG antibody was reverse transcribed using the Superscript III First-Strand Synthesis system and qPCR was performed using iTaq Universal SYBR Green Supermix with the ABI 7900HT.Percent of input was calculated against input. For RT-qPCR primers and antibodies see Table S1.

### CUT&Tag library preparation and data processing

CUT&Tag libraries were prepared using the CUT&Tag-IT Assay Kit (Active Motif, 53160) following to the manufacturer’s protocol. 10^6^ of MEL cells were collected for each biological replicate and two replicates were prepared. The MEL cells were bound to Concanavalin A Beads and Incubated with 1:50 rabbit polyclonal Phospho-Rpb1 CTD Ser5 and Ser2 antibody (Cell Signaling 13523 and 13499). Guinea pig a-rabbit antibody was used at 1:100 dilution as secondary antibody. Tagmentation was performed using pA-Tn5 Transposomes at 37°C for 60 minutes. DNA was purified by DNA Purification Column, then universal i5 primers and uniquely barcoded i7 primers were added to the DNA with 14 cycles of PCR. Individual libraries were purified with SPRI beads and eluted with 20 µL DNA Purification Buffer. The libraries were sequenced on a MiSeq. Antibodies are listed in Table S1.

### CUT&Tag data analysis

Raw fastq pairs were preprocessed and removed adapter and low-quality sequences using the cutadapt program (v2.7) ^71^. Preprocessed reads were aligned to the mouse genome available at Gencode M18 ^72^ using Bowtie2 (v2.3.5) ^73^ with the settings for Cut&Tag ^74^. Final reads were retained after removing non-uniquely mapped reads using samtools (v1.9) with “-q 20” ^75^ and duplicated reads using picard (v2.21.4). Peaks were called using MACS2 (version 2.2.7.1) ^76,77^. Browser track files were generated using the deepTools (v3.3.1) ^78^.

## QUANTIFICATION AND STATISTICAL ANALYSIS

### RNA Polymerase II (Pol2) pausing index at *Myb* locus

For Pol2 abundance over specific regions and Pol2 pausing analysis, bam files were filtered for properly paired reads only with samtools view, and then converted to bedpe format using bedtools bamtobed. The subset of fragments that were shorter than 1000 bp and mapped to chromosomes chr 1-19, X, Y was extracted to generate a bed file of fragments. Subsets of bed files were generated based on the desired size ranges (total, <120 bp, 120-270 bp, >270 bp). Fragments over genomic regions of interest were counted with bedtools intersect. The Pol2 pausing index was calculated as the ratio of the fragments per million (FPM) in the first exon of *Myb* over the rest of the gene body.

### Statistical analysis

As indicated in the figure legends, data values reported in the figures are the mean and standard error of the mean (SEM). GraphPad Prism 8.0 (GraphPad Software) and Excel (Microsoft) were used to perform the statistical analyses. Unpaired Student’s *t*-test was used for significance test. (*) P < 0.05, (**) P < 0.01, (***) P < 0.001.

## Notes

### Competing Interest Statement

The authors have declared no competing interest.

### Summary of Updates

The method part was edited.

## REFERENCES

1. Cabili, M.N., Trapnell, C., Goff, L., Koziol, M., Tazon-Vega, B., Regev, A., and Rinn, J.L. (2011). Integrative annotation of human large intergenic noncoding RNAs reveals global properties and specific subclasses. Genes Dev 25, 1915–1927. gad.17446611 [pii];10.1101/gad.17446611 [doi].

2. Derrien, T., Johnson, R., Bussotti, G., Tanzer, A., Djebali, S., Tilgner, H., Guernec, G., Martin, D., Merkel, A., Knowles, D.G., et al. (2012). The GENCODE v7 catalog of human long noncoding RNAs: analysis of their gene structure, evolution, and expression. Genome. Res 22, 1775–1789. 22/9/1775 [pii];10.1101/gr.132159.111 [doi].

3. Djebali, S., Davis, C.A., Merkel, A., Dobin, A., Lassmann, T., Mortazavi, A., Tanzer, A., Lagarde, J., Lin, W., Schlesinger, F., et al. (2012). Landscape of transcription in human cells. Nature 489, 101–108. nature11233 [pii];10.1038/nature11233 [doi].

4. Hu, W., Alvarez-Dominguez, J.R., and Lodish, H.F. (2012). Regulation of mammalian cell differentiation by long non-coding RNAs. EMBO Rep 13, 971–983. embor2012145 [pii];10.1038/embor.2012.145 [doi].

5. Kim, T.K., and Shiekhattar, R. (2016). Diverse regulatory interactions of long noncoding RNAs. Curr. Opin. Genet. Dev 36, 73–82. S0959-437X(16)30023-5 [pii];10.1016/j.gde.2016.03.014 [doi].

6. Gil, N., and Ulitsky, I. (2020). Regulation of gene expression by cis-acting long non-coding RNAs. Nat Rev Genet 21, 102–117. 10.1038/s41576-019-0184-5.

7. Statello, L., Guo, C.J., Chen, L.L., and Huarte, M. (2021). Gene regulation by long non-coding RNAs and its biological functions. Nat Rev Mol Cell Biol 22, 96–118. 10.1038/s41580-020-00315-9.

8. Herman, A.B., Tsitsipatis, D., and Gorospe, M. (2022). Integrated lncRNA function upon genomic and epigenomic regulation. Mol Cell 82, 2252–2266. 10.1016/j.molcel.2022.05.027.

9. Arnold, P.R., Wells, A.D., and Li, X.C. (2019). Diversity and Emerging Roles of Enhancer RNA in Regulation of Gene Expression and Cell Fate. Front Cell Dev Biol 7, 377. 10.3389/fcell.2019.00377.

10. Sartorelli, V., and Lauberth, S.M. (2020). Enhancer RNAs are an important regulatory layer of the epigenome. Nat Struct Mol Biol 27, 521–528. 10.1038/s41594-020-0446-0.

11. Hou, T.Y., and Kraus, W.L. (2021). Spirits in the Material World: Enhancer RNAs in Transcriptional Regulation. Trends Biochem Sci 46, 138–153. 10.1016/j.tibs.2020.08.007.

12. Plank, J.L., and Dean, A. (2014). Enhancer Function: Mechanistic and Genome-Wide Insights Come Together. Mol. Cell 55, 5–14. S1097-2765(14)00523-1 [pii];10.1016/j.molcel.2014.06.015 [doi].

13. Levine, M., Cattoglio, C., and Tjian, R. (2014). Looping back to leap forward: transcription enters a new era. Cell 157, 13–25. S0092-8674(14)00201-3 [pii];10.1016/j.cell.2014.02.009 [doi].

14. Furlong, E.E.M., and Levine, M. (2018). Developmental enhancers and chromosome topology. Science 361, 1341–1345. 10.1126/science.aau0320.

15. Panigrahi, A., and O’Malley, B.W. (2021). Mechanisms of enhancer action: the known and the unknown. Genome Biol 22, 108. 10.1186/s13059-021-02322-1.

16. Kim, J., and Dean, A. (2021). Enhancers navigate the three-dimensional genome to direct cell fate decisions. Curr Opin Struct Biol 71, 101–109. 10.1016/j.sbi.2021.06.005.

17. Kim, S., and Wysocka, J. (2023). Deciphering the multi-scale, quantitative cis-regulatory code. Mol Cell 83, 373–392. 10.1016/j.molcel.2022.12.032.

18. Dou, Y., Milne, T.A., Ruthenburg, A.J., Lee, S., Lee, J.W., Verdine, G.L., Allis, C.D., and Roeder, R.G. (2006). Regulation of MLL1 H3K4 methyltransferase activity by its core components. Nat Struct Mol Biol 13, 713–719. 10.1038/nsmb1128.

19. Cenik, B.K., and Shilatifard, A. (2021). COMPASS and SWI/SNF complexes in development and disease. Nat Rev Genet 22, 38–58. 10.1038/s41576-020-0278-0.

20. Milne, T.A., Dou, Y., Martin, M.E., Brock, H.W., Roeder, R.G., and Hess, J.L. (2005). MLL associates specifically with a subset of transcriptionally active target genes. Proc Natl Acad Sci U S A 102, 14765–14770. 10.1073/pnas.0503630102.

21. Birke, M., Schreiner, S., Garcia-Cuellar, M.P., Mahr, K., Titgemeyer, F., and Slany, R.K. (2002). The MT domain of the proto-oncoprotein MLL binds to CpG-containing DNA and discriminates against methylation. Nucleic Acids Res 30, 958–965. 10.1093/nar/30.4.958.

22. Xu, J., Li, L., Xiong, J., denDekker, A., Ye, A., Karatas, H., Liu, L., Wang, H., Qin, Z.S., Wang, S., and Dou, Y. (2016). MLL1 and MLL1 fusion proteins have distinct functions in regulating leukemic transcription program. Cell Discov 2, 16008. 10.1038/celldisc.2016.8.

23. Wang, K.C., Yang, Y.W., Liu, B., Sanyal, A., Corces-Zimmerman, R., Chen, Y., Lajoie, B.R., Protacio, A., Flynn, R.A., Gupta, R.A., et al. (2011). A long noncoding RNA maintains active chromatin to coordinate homeotic gene expression. Nature 472, 120–124. 10.1038/nature09819.

24. Deng, C., Li, Y., Zhou, L., Cho, J., Patel, B., Terada, N., Li, Y., Bungert, J., Qiu, Y., and Huang, S. (2016). HoxBlinc RNA Recruits Set1/MLL Complexes to Activate Hox Gene Expression Patterns and Mesoderm Lineage Development. Cell Rep 14, 103–114. 10.1016/j.celrep.2015.12.007.

25. Wang, X., Angelis, N., and Thein, S.L. (2018). MYB - A regulatory factor in hematopoiesis. Gene 665, 6–17. S0378-1119(18)30439-6 [pii];10.1016/j.gene.2018.04.065 [doi].

26. Jiang, J., Best, S., Menzel, S., Silver, N., Lai, M.I., Surdulescu, G.L., Spector, T.D., and Thein, S.L. (2006). cMYB is involved in the regulation of fetal hemoglobin production in adults. Blood 108, 1077–1083. 108/3/1077 [pii];10.1182/blood-2006-01-008912 [doi].

27. Sankaran, V.G., Menne, T.F., Scepanovic, D., Vergilio, J.A., Ji, P., Kim, J., Thiru, P., Orkin, S.H., Lander, E.S., and Lodish, H.F. (2011). MicroRNA-15a and −16-1 act via MYB to elevate fetal hemoglobin expression in human trisomy 13. Proc Natl Acad Sci U. S. A 108, 1519–1524. 1018384108 [pii];10.1073/pnas.1018384108 [doi].

28. Stadhouders, R., Thongjuea, S., Andrieu-Soler, C., Palstra, R.J., Bryne, J.C., van den Heuvel, A., Stevens, M., de, B.E., Kockx, C., van der Sloot, A., et al. (2012). Dynamic long-range chromatin interactions control Myb proto-oncogene transcription during erythroid development. EMBO J 31, 986–999. emboj2011450 [pii];10.1038/emboj.2011.450 [doi].

29. Stadhouders, R., Aktuna, S., Thongjuea, S., Aghajanirefah, A., Pourfarzad, F., van, I.W., Lenhard, B., Rooks, H., Best, S., Menzel, S., et al. (2014). HBS1L-MYB intergenic variants modulate fetal hemoglobin via long-range MYB enhancers. J. Clin. Invest 124, 1699–1710. 71520 [pii];10.1172/JCI71520 [doi].

30. Wahlberg, K., Jiang, J., Rooks, H., Jawaid, K., Matsuda, F., Yamaguchi, M., Lathrop, M., Thein, S.L., and Best, S. (2009). The HBS1L-MYB intergenic interval associated with elevated HbF levels shows characteristics of a distal regulatory region in erythroid cells. Blood 114, 1254–1262. blood-2009-03-210146 [pii];10.1182/blood-2009-03-210146 [doi].

31. Stadhouders, R., Cico, A., Stephen, T., Thongjuea, S., Kolovos, P., Baymaz, H.I., Yu, X., Demmers, J., Bezstarosti, K., Maas, A., et al. (2015). Control of developmentally primed erythroid genes by combinatorial co-repressor actions. Nat Commun 6, 8893. 10.1038/ncomms9893.

32. Song, S.-H., Hou, C., and Dean, A. (2007). A positive role for NLI/Ldb1 in long-range β-globin locus control region function. Mol. Cell 28, 810–822.

33. Alvarez-Dominguez, J.R., Hu, W., Yuan, B., Shi, J., Park, S.S., Gromatzky, A.A., van, O.A., and Lodish, H.F. (2014). Global discovery of erythroid long noncoding RNAs reveals novel regulators of red cell maturation. Blood 123, 570–581. blood-2013-10-530683 [pii];10.1182/blood-2013-10-530683 [doi].

34. Ivaldi, M.S., Diaz, L.F., Chakalova, L., Lee, J., Krivega, I., and Dean, A. (2018). Fetal gamma-globin genes are regulated by the BGLT3 long noncoding RNA locus. Blood 132, 1963–1973. blood-2018-07-862003 [pii];10.1182/blood-2018-07-862003 [doi].

35. Paralkar, V.R., Mishra, T., Luan, J., Yao, Y., Kossenkov, A.V., Anderson, S.M., Dunagin, M., Pimkin, M., Gore, M., Sun, D., et al. (2014). Lineage and species-specific long noncoding RNAs during erythro-megakaryocytic development. Blood 123, 1927–1937. blood-2013-12-544494 [pii];10.1182/blood-2013-12-544494 [doi].

36. Morrison, T.A., Wilcox, I., Luo, H.Y., Farrell, J.J., Kurita, R., Nakamura, Y., Murphy, G.J., Cui, S., Steinberg, M.H., and Chui, D.H.K. (2018). A long noncoding RNA from the HBS1L-MYB intergenic region on chr6q23 regulates human fetal hemoglobin expression. Blood Cells Mol. Dis 69, 1–9. S1079-9796(17)30440-0 [pii];10.1016/j.bcmd.2017.11.003 [doi].

37. Hu, W., Yuan, B., Flygare, J., and Lodish, H.F. (2011). Long noncoding RNA-mediated anti-apoptotic activity in murine erythroid terminal differentiation. Genes Dev 25, 2573–2578. gad.178780.111 [pii];10.1101/gad.178780.111 [doi].

38. Gurumurthy, A., Yu, D.T., Stees, J.R., Chamales, P., Gavrilova, E., Wassel, P., Li, L., Stribling, D., Chen, J., Brackett, M., et al. (2021). Super-enhancer mediated regulation of adult beta-globin gene expression: the role of eRNA and Integrator. Nucleic Acids Res 49, 1383–1396. 10.1093/nar/gkab002.

39. Wang, L., Park, H.J., Dasari, S., Wang, S., Kocher, J.P., and Li, W. (2013). CPAT: Coding-Potential Assessment Tool using an alignment-free logistic regression model. Nucleic Acids Res 41, e74. 10.1093/nar/gkt006.

40. Li, W., Notani, D., and Rosenfeld, M.G. (2016). Enhancers as non-coding RNA transcription units: recent insights and future perspectives. Nat. Rev. Genet 17, 207–223. nrg.2016.4 [pii];10.1038/nrg.2016.4 [doi].

41. Tober, J., McGrath, K.E., and Palis, J. (2008). Primitive erythropoiesis and megakaryopoiesis in the yolk sac are independent of c-myb. Blood 111, 2636–2639. blood-2007-11-124685 [pii];10.1182/blood-2007-11-124685 [doi].

42. Castro-Mondragon, J.A., Riudavets-Puig, R., Rauluseviciute, I., Lemma, R.B., Turchi, L., Blanc-Mathieu, R., Lucas, J., Boddie, P., Khan, A., Manosalva Perez, N., et al. (2022). JASPAR 2022: the 9th release of the open-access database of transcription factor binding profiles. Nucleic Acids Res 50, D165–D173. 10.1093/nar/gkab1113.

43. Henriques, T., Scruggs, B.S., Inouye, M.O., Muse, G.W., Williams, L.H., Burkholder, A.B., Lavender, C.A., Fargo, D.C., and Adelman, K. (2018). Widespread transcriptional pausing and elongation control at enhancers. Genes Dev 32, 26–41. 10.1101/gad.309351.117.

44. Mucenski, M.L., McLain, K., Kier, A.B., Swerdlow, S.H., Schreiner, C.M., Miller, T.A., Pietryga, D.W., Scott, W.J., Jr., and Potter, S.S. (1991). A functional c-myb gene is required for normal murine fetal hepatic hematopoiesis. Cell 65, 677–689. 10.1016/0092-8674(91)90099-k.

45. Bauer, D.E., and Orkin, S.H. (2011). Update on fetal hemoglobin gene regulation in hemoglobinopathies. Curr. Opin. Pediatr 23, 1–8. 10.1097/MOP.0b013e3283420fd0 [doi].

46. Bianchi, E., Zini, R., Salati, S., Tenedini, E., Norfo, R., Tagliafico, E., Manfredini, R., and Ferrari, S. (2010). c-myb supports erythropoiesis through the transactivation of KLF1 and LMO2 expression. Blood 116, e99–110. 10.1182/blood-2009-08-238311.

47. Lee, J., Krivega, I., Dale, R.K., and Dean, A. (2017). The LDB1 Complex Co-opts CTCF for Erythroid Lineage-Specific Long-Range Enhancer Interactions. Cell Rep 19, 2490–2502. S2211-1247(17)30745-3 [pii];10.1016/j.celrep.2017.05.072 [doi].

48. Nguyen, N., Gudmundsson, K.O., Soltis, A.R., Oakley, K., Roy, K.R., Han, Y., Gurnari, C., Maciejewski, J.P., Crouch, G., Ernst, P., et al. (2022). Recruitment of MLL1 complex is essential for SETBP1 to induce myeloid transformation. iScience 25, 103679. 10.1016/j.isci.2021.103679.

49. Mishra, B.P., Zaffuto, K.M., Artinger, E.L., Org, T., Mikkola, H.K., Cheng, C., Djabali, M., and Ernst, P. (2014). The histone methyltransferase activity of MLL1 is dispensable for hematopoiesis and leukemogenesis. Cell Rep 7, 1239–1247. 10.1016/j.celrep.2014.04.015.

50. Fujinaga, K., Huang, F., and Peterlin, B.M. (2023). P-TEFb: The master regulator of transcription elongation. Mol Cell 83, 393–403. 10.1016/j.molcel.2022.12.006.

51. Mitra, P., Pereira, L.A., Drabsch, Y., Ramsay, R.G., and Gonda, T.J. (2012). Estrogen receptor-alpha recruits P-TEFb to overcome transcriptional pausing in intron 1 of the MYB gene. Nucleic Acids Res 40, 5988–6000. 10.1093/nar/gks286.

52. Miura, M., and Chen, H. (2020). CUT&RUN detects distinct DNA footprints of RNA polymerase II near the transcription start sites. Chromosome Res 28, 381–393. 10.1007/s10577-020-09643-0.

53. Oksuz, O., Henninger, J.E., Warneford-Thomson, R., Zheng, M.M., Erb, H., Vancura, A., Overholt, K.J., Hawken, S.W., Banani, S.F., Lauman, R., et al. (2023). Transcription factors interact with RNA to regulate genes. Mol Cell 83, 2449–2463 e2413. 10.1016/j.molcel.2023.06.012.

54. Bassett, A.R., Akhtar, A., Barlow, D.P., Bird, A.P., Brockdorff, N., Duboule, D., Ephrussi, A., Ferguson-Smith, A.C., Gingeras, T.R., Haerty, W., et al. (2014). Considerations when investigating lncRNA function in vivo. Elife 3, e03058. 10.7554/eLife.03058 [doi].

55. Oudelaar, A.M., Davies, J.O.J., Hanssen, L.L.P., Telenius, J.M., Schwessinger, R., Liu, Y., Brown, J.M., Downes, D.J., Chiariello, A.M., Bianco, S., et al. (2018). Single-allele chromatin interactions identify regulatory hubs in dynamic compartmentalized domains. Nat Genet 50, 1744–1751. 10.1038/s41588-018-0253-2.

56. Proudhon, C., Snetkova, V., Raviram, R., Lobry, C., Badri, S., Jiang, T., Hao, B., Trimarchi, T., Kluger, Y., Aifantis, I., et al. (2016). Active and Inactive Enhancers Cooperate to Exert Localized and Long-Range Control of Gene Regulation. Cell Rep 15, 2159–2169. 10.1016/j.celrep.2016.04.087.

57. Jiang, T., Raviram, R., Snetkova, V., Rocha, P.P., Proudhon, C., Badri, S., Bonneau, R., Skok, J.A., and Kluger, Y. (2016). Identification of multi-loci hubs from 4C-seq demonstrates the functional importance of simultaneous interactions. Nucleic Acids Res 44, 8714–8725. 10.1093/nar/gkw568.

58. Fudenberg, G., Imakaev, M., Lu, C., Goloborodko, A., Abdennur, N., and Mirny, L.A. (2016). Formation of Chromosomal Domains by Loop Extrusion. Cell Rep 15, 2038–2049. S2211-1247(16)30530-7 [pii];10.1016/j.celrep.2016.04.085 [doi].

59. Sanborn, A.L., Rao, S.S., Huang, S.C., Durand, N.C., Huntley, M.H., Jewett, A.I., Bochkov, I.D., Chinnappan, D., Cutkosky, A., Li, J., et al. (2015). Chromatin extrusion explains key features of loop and domain formation in wild-type and engineered genomes. Proc. Natl. Acad. Sci. U. S. A 112, E6456–E6465. 1518552112 [pii];10.1073/pnas.1518552112 [doi].

60. Krivega, I., Dale, R.K., and Dean, A. (2014). Role of LDB1 in the transition from chromatin looping to transcription activation. Genes Dev 28, 1278–1290. gad.239749.114 [pii];10.1101/gad.239749.114 [doi].

61. Palstra, R.J., Simonis, M., Klous, P., Brasset, E., Eijkelkamp, B., and de, L.W. (2008). Maintenance of Long-Range DNA Interactions after Inhibition of Ongoing RNA Polymerase II Transcription. PLoS. ONE 3, e1661.

62. Crump, N.T., Ballabio, E., Godfrey, L., Thorne, R., Repapi, E., Kerry, J., Tapia, M., Hua, P., Lagerholm, C., Filippakopoulos, P., et al. (2021). BET inhibition disrupts transcription but retains enhancer-promoter contact. Nat Commun 12, 223. 10.1038/s41467-020-20400-z.

63. Murphy, Z.C., Murphy, K., Myers, J., Getman, M., Couch, T., Schulz, V.P., Lezon-Geyda, K., Palumbo, C., Yan, H., Mohandas, N., et al. (2021). Regulation of RNA polymerase II activity is essential for terminal erythroid maturation. Blood 138, 1740–1756. 10.1182/blood.2020009903.

64. Ghavi-Helm, Y., Klein, F.A., Pakozdi, T., Ciglar, L., Noordermeer, D., Huber, W., and Furlong, E.E. (2014). Enhancer loops appear stable during development and are associated with paused polymerase. Nature 512, 96–100. nature13417 [pii];10.1038/nature13417 [doi].

65. Gressel, S., Schwalb, B., Decker, T.M., Qin, W., Leonhardt, H., Eick, D., and Cramer, P. (2017). CDK9-dependent RNA polymerase II pausing controls transcription initiation. Elife 6. 10.7554/eLife.29736.

66. Rahl, P.B., Lin, C.Y., Seila, A.C., Flynn, R.A., McCuine, S., Burge, C.B., Sharp, P.A., and Young, R.A. (2010). c-Myc regulates transcriptional pause release. Cell 141, 432–445. 10.1016/j.cell.2010.03.030.

67. Wang, H., Fan, Z., Shliaha, P.V., Miele, M., Hendrickson, R.C., Jiang, X., and Helin, K. (2023). H3K4me3 regulates RNA polymerase II promoter-proximal pause-release. Nature 615, 339–348. 10.1038/s41586-023-05780-8.

68. Farrell, J.J., Sherva, R.M., Chen, Z.Y., Luo, H.Y., Chu, B.F., Ha, S.Y., Li, C.K., Lee, A.C., Li, R.C., Li, C.K., et al. (2011). A 3-bp deletion in the HBS1L-MYB intergenic region on chromosome 6q23 is associated with HbF expression. Blood 117, 4935–4945. 10.1182/blood-2010-11-317081.

69. Hezroni, H., Koppstein, D., Schwartz, M.G., Avrutin, A., Bartel, D.P., and Ulitsky, I. (2015). Principles of long noncoding RNA evolution derived from direct comparison of transcriptomes in 17 species. Cell Rep 11, 1110–1122. 10.1016/j.celrep.2015.04.023.

70. Ran, F.A., Hsu, P.D., Wright, J., Agarwala, V., Scott, D.A., and Zhang, F. (2013). Genome engineering using the CRISPR-Cas9 system. Nat. Protoc 8, 2281–2308. nprot.2013.143 [pii];10.1038/nprot.2013.143 [doi].

71. Martin, M. (2011). Cutadapt removes adapter sequences from high-throughput sequencing reads. 2011 17, 3. 10.14806/ej.17.1.200.

72. Frankish, A., Carbonell-Sala, S., Diekhans, M., Jungreis, I., Loveland, J.E., Mudge, J.M., Sisu, C., Wright, J.C., Arnan, C., Barnes, I., et al. (2023). GENCODE: reference annotation for the human and mouse genomes in 2023. Nucleic Acids Res 51, D942–D949. 10.1093/nar/gkac1071.

73. Langmead, B., and Salzberg, S.L. (2012). Fast gapped-read alignment with Bowtie 2. Nat Methods 9, 357–359. 10.1038/nmeth.1923.

74. Kaya-Okur, H.S., Wu, S.J., Codomo, C.A., Pledger, E.S., Bryson, T.D., Henikoff, J.G., Ahmad, K., and Henikoff, S. (2019). CUT&Tag for efficient epigenomic profiling of small samples and single cells. Nat Commun 10, 1930. 10.1038/s41467-019-09982-5.

75. Danecek, P., Bonfield, J.K., Liddle, J., Marshall, J., Ohan, V., Pollard, M.O., Whitwham, A., Keane, T., McCarthy, S.A., Davies, R.M., and Li, H. (2021). Twelve years of SAMtools and BCFtools. Gigascience 10. 10.1093/gigascience/giab008.

76. Zhang, Y., Liu, T., Meyer, C.A., Eeckhoute, J., Johnson, D.S., Bernstein, B.E., Nusbaum, C., Myers, R.M., Brown, M., Li, W., and Liu, X.S. (2008). Model-based analysis of ChIP-Seq (MACS). Genome Biol 9, R137. 10.1186/gb-2008-9-9-r137.

77. Liu, T. (2014). Use model-based Analysis of ChIP-Seq (MACS) to analyze short reads generated by sequencing protein-DNA interactions in embryonic stem cells. Methods Mol Biol 1150, 81–95. 10.1007/978-1-4939-0512-6_4.

78. Ramirez, F., Dundar, F., Diehl, S., Gruning, B.A., and Manke, T. (2014). deepTools: a flexible platform for exploring deep-sequencing data. Nucleic Acids Res 42, W187–191. 10.1093/nar/gku365.

79. Vakoc, C.R., Mandat, S.A., Olenchock, B.A., and Blobel, G.A. (2005). Histone H3 lysine 9 methylation and HP1gamma are associated with transcription elongation through mammalian chromatin. Mol Cell 19, 381–391. 10.1016/j.molcel.2005.06.011.

80. Doench, J.G., Fusi, N., Sullender, M., Hegde, M., Vaimberg, E.W., Donovan, K.F., Smith, I., Tothova, Z., Wilen, C., Orchard, R., et al. (2016). Optimized sgRNA design to maximize activity and minimize off-target effects of CRISPR-Cas9. Nat Biotechnol 34, 184–191. 10.1038/nbt.3437.

