## Supplemental data for "An enhancer RNA recruits MLL1 to regulate transcription of *Myb*"

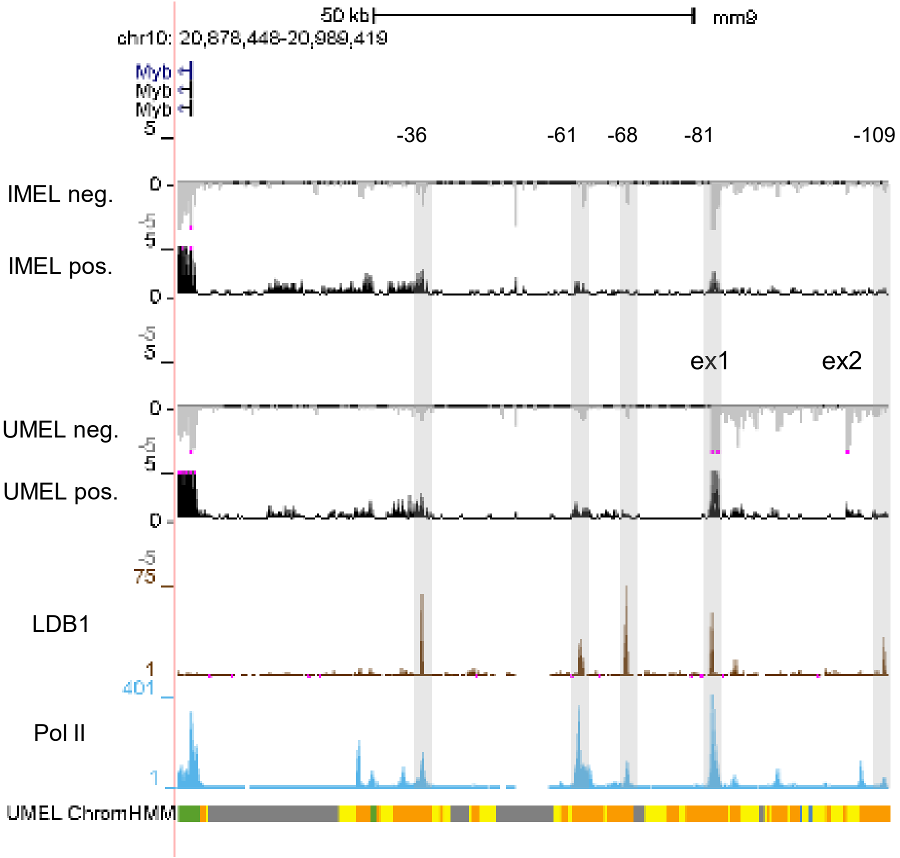


**Figure S1. RNA-seq results for the *Myb*-Hbsl1 intergenic region**. Signal tracks for RNA-seq of induced (IMEL) and uninduced (UMEL) MEL cells from ENCODE are shown together with signal tracks for LDB1 and Pol II. Related to Figure 1.

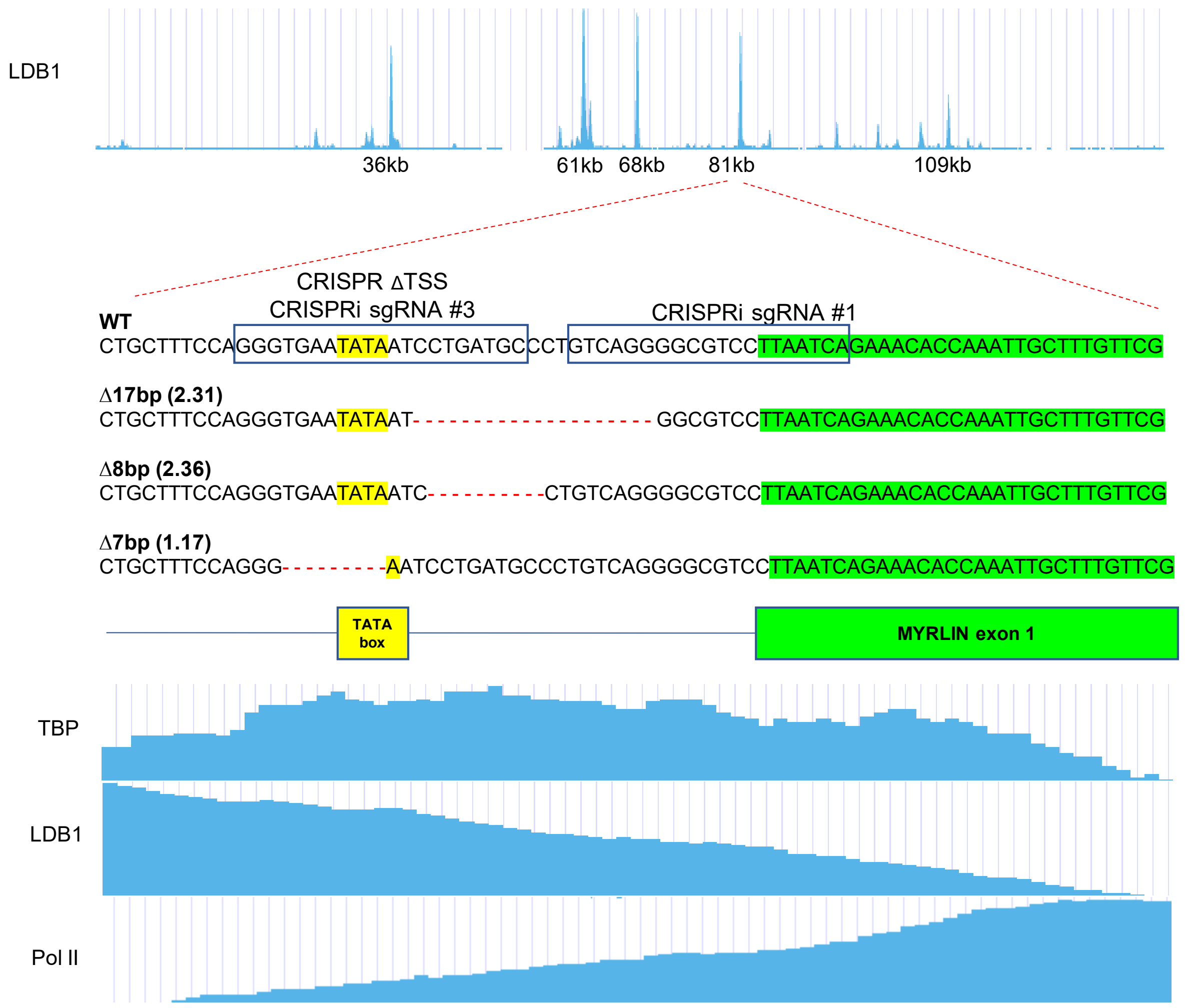


**Figure S2. Schematic of the *Myb-Hbsl1* intergenic region.** The zoomed in portion at the -81kb enhancer shows the gRNA used for targeting the *Myrlin* TSS, the resulting three CRISPR-Cas9 deletions from Figure 2, the TATA box and the transcription start site for *Myrlin*. Below are signal tracks for TBP, LDB1 and Pol II from ENCODE. The locations of gRNAs used for CRISPR targeting of dCAS9-KRAB are also indicated. Related to Figure 2.

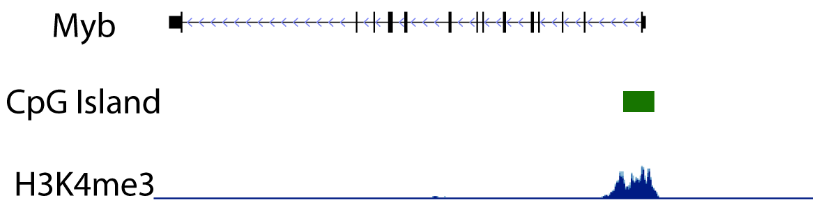


**Figure S3. Encode genome browser view of the CpG island in the *Myb* promoter.** The *Myb* promoter and first exon/intron contain a CpG island, which is highly enriched for H3K4me3 when *Myb* is actively transcribed. Related to Figure 5.

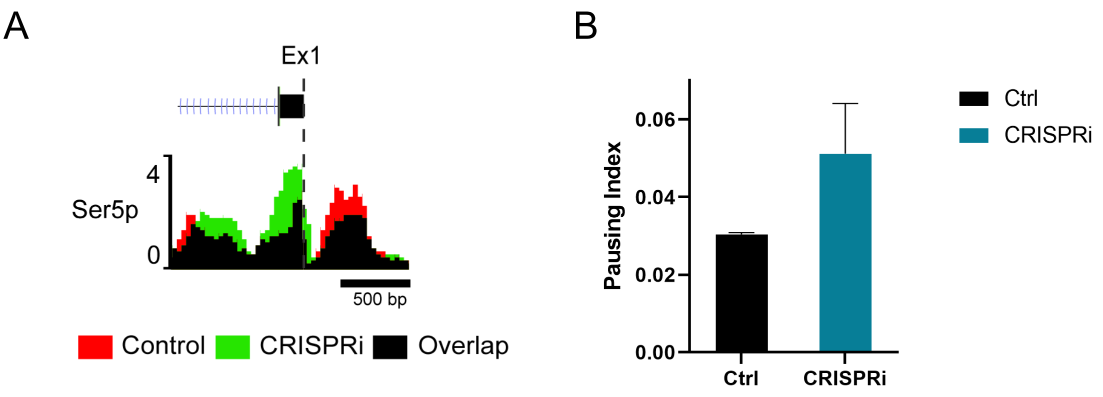


**Figure S4**. **Pol II Ser5P occupancy in the *Myb* promoter.** (A) Analysis of the Pol II Ser5P CUT & TAG fragments displaying Pol II Ser5 occupancy. (B) Pausing index calculated for Pol II Ser5 across *Myb*. Error bars indicate SEM of 2 independent experiments. Related to Figure 6.

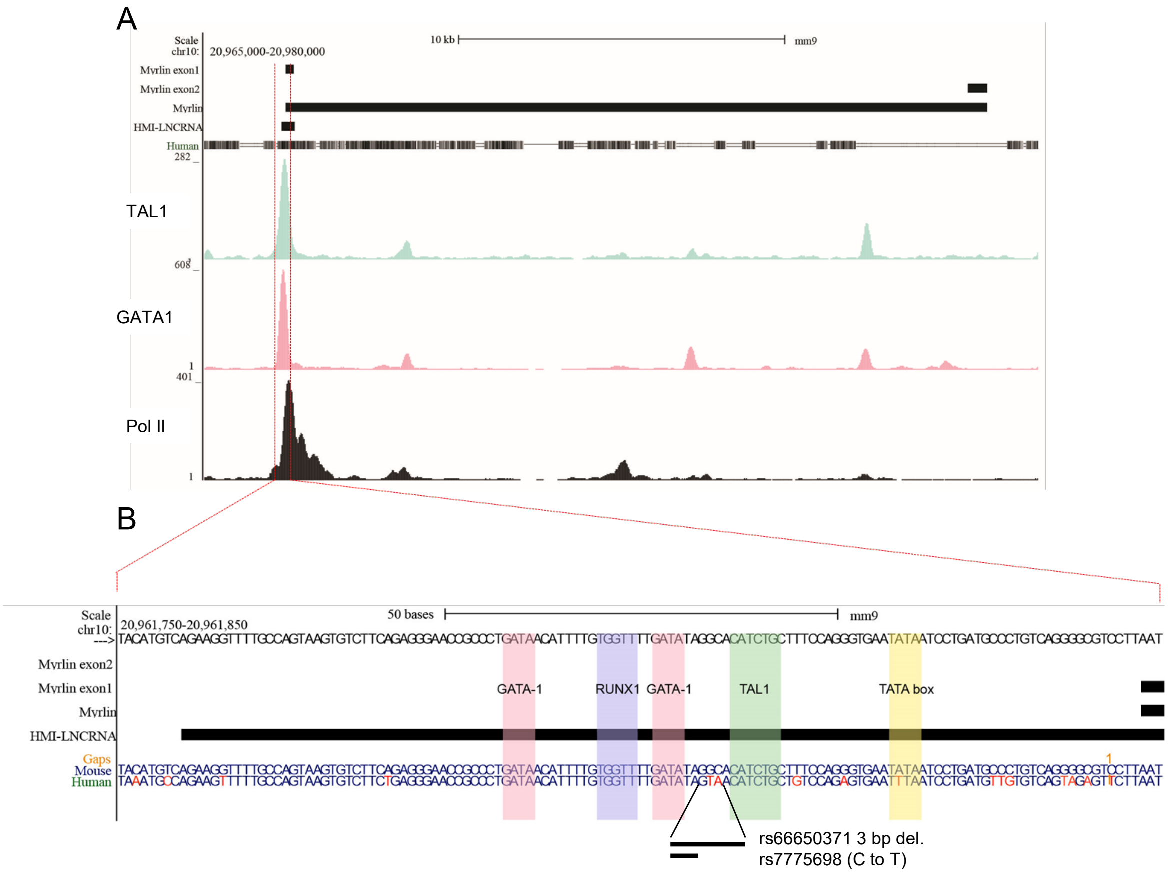


**Figure S5**. **Human and mouse *Myb* enhancer transcripts.** (A) Illustration of genomic region between chr10: 20,965,000 – 20,980,000 (mm9), showing BLATN result for human HMI-LNCRNA lncRNA and a 378 bp homology patch (see text) shared with the mouse *Myrlin* locus. Below are ENCODE ChIP-seq signal tracks for TAL1, GATA1, and Pol2 at the -81kb intergenic enhancer. (B) Chr10: 20,961,750 – 20,961,850 (mm9) zoom-in snapshot (dashed red lines) including HMI-LNCRNA DNA homology patch and highlighting GATA1, RUNX1, and TAL1 motifs conserved between mouse and human. The TATA box motif (yellow) appears only in the mouse reference genome. Below are the mouse and human DNA sequence showing nucleotide differences (red) with human single nucleotide polymorphisms (SNPs) rs66650371 and rs7775698 that are among the SNPs in the MYB locus that are linked to elevated fetal hemoglobin in humans.

**Table S1. List of primers and oligos. Related to STAR Methods.**

| | Sequence 5'-3' |
|---|---|
| CRISPR-gRNA | |
| gRNA1 -81kb TSS SE | CACCGTCAGGGGCGTCCTTAATCA |
| gRNA1 -81kb TSS AE | AAACTGATTAAGGACGCCCCTGAC |
| gRNA2 -81kb TSS SE | CACCGGACGCCCCTGACAGGGCATC |
| gRNA2 -81kb TSS AE | AAACGATGCCCTGTCAGGGGCGTCC |
| gRNA3 -81kb TSS SE | CACCGCATCAGGATTATATTCACCC |
| gRNA3 -81kb TSS AE | AAACGGGTGAATATAATCCTGATGC |
| RACE | |
| 3' RACE outer | GAGTTTCACAACCACCTTCCAA |
| 3' RACE inner | GAGAACAACAGTGGATGGGC |
| 5' RACE outer | TAGCTGAGGGAACAAACGGAG |
| 5' RACE inner | CTCTGGCATAACTGGAGTAAATTA |
| PCR | |
| Myrlin exon 1 F | TTAATCAGAAACACCAAATTGCTT |
| Myrlin exon 1 F | CTGTAGCTGAGGGAACAAACG |
| Myrlin exon 2 F | ATCAGGATACAGCTCTCAGCTACT |
| Myrlin exon 2 R | CGAGGCAATCTCTTGGTTTTA |
| RT & ChIP qPCR | |
| Actin exon F | ACCTTCTACAATGAGCTGCG |
| Actin exon R | CTGGATGGCTACGTACATGG |
| Actin intron F | GGGGCTCCACTTAGACCTAC |
| Actin intron R | GGCTCGTGTGACAAAGCTAA |
| Myrlin exon 1 F | AAATTGCTTTGTTCGCCTGC |
| Myrlin exon 1 R | CGAGTACCGCTTTCACAGTT |
| Myrlin exon 1/2 junction F | GTTCCGGCTCCGTTTGTT |
| Myrlin exon 1/2 junction R | CATCATTATGGTGGGGAACAT |
| Myrlin exon 2 F | TCAGCTACTGTTCCAGTGCC |
| Myrlin exon 2 R | GGGCTTACAGTTCCAGAGGA |
| Malat1 F | CAGGAGTGAGGCTTGTGGTA |
| Malat1 R | TGAATGGTAGAGCCAGCACA |
| Myb prom F | TCTTTGTTTGATGGCATCTGTT |
| Myb prom R | AAAGGGGAGGAGAAGGAGGT |
| -36kb F | TCACTTCCTTCCTGTCTCG |
| -36kb R | GTCTGGTGGCGATGACTTA |
| -61kb F | GTTGGGCAAAGATACTGGAT |
| -61kb R | GTTGGGCAAAGATACTGGAT |
| -68kb F | ATCCAAGTGACGGTGACA |
| -68kb R | GCATCCTGATTGTGCTAACT |
| -81kb F | CCCAAGTGAGAGAAATGTTGAA |
| -81kb R | TGTTATCAGGGCGGTTCC |

| | |
|---|---|
| -109kb F | CTCCAACATCAGCCGACT |
| -109kb R | ATTGTGAGGTGAGTGCCC |
| Myb exon 2 F | TTTCCAGATTTGGGCAGCAG |
| Myb exon 2 R | TGCAGCATCTACAGTAGCGA |
| Myb exon 3 F | GGTCCAGGGACCTTTGATGA |
| Myb exon 3 R | GACAGATGTGCAGTGCCAA |
| Necdin F | AGCTCATGTGGTACGTGTTGGT |
| Necdin R | GCTGCCCATGACCTCTTTCA |
| HS2 F | ATGTTAGTGTCAGCATATTACCGATGT |
| HS2 R | AATGCAGAGGCTCGTTAATGTG |
| Myb int1 F | GAACTGCAAACCAGCCTCTT |
| Myb int1 R | GGGATGTTTCATCCTGAGAGC |
| Hbb-bh1 F | CCTGGCCATCATGGGAAAC |
| Hbb-bh1 R | CCCCAAGCCCAAGGATGT |
| Hbb-b1 F | CACATGCAGCTTGTCACAGT |
| Hbb-b1 R | GTCAAAGCCCATGGCAAGAA |
| Hbb-y F | GCTTGTCACAGTGCAGTTCA |
| Hbb-y R | CGATGGCCTGAATCACTTGG |
| Chromatin Capture 3C Taqman qPCR | |
| Myb prom universal F | AGTTCAAGACTTGTGCTGACTG |
| Myb prom universal Probe | 5’-FAM-AACCCCTAAAGCACTTGGGACACTCACAC-3’-BHQ1 |
| 20898686 R | CCTAGGTTCTCCTCTCCTACCA |
| 20904395 R | GACAATTTGACATGAATTGCAAGC |
| 20909424 R | CAAGAACCAAGACGCCTCAG |
| 20915439 R | AGTAAATCTTGCTGCCCTCAAG |
| 20923231 R | TGCCTTGGGCAGTTCTATAGAG |
| 20939397 R | TCCTTACTTCTGCTCTCAAACAC |
| 20944478 R | CTTTGTAGGTCACTTTCTCCAGC |
| 20947418 R | AGCAGGCTATTGTGAAAAGAGG |
| 20950525 R | ACACATAGGCACAGAGGAAAGTT |
| 20957119 R | GCAACCTTTTCACTGGCAGAAAT |
| 20974886 R | GAAGACCACTTAGGTAAACACTTTG |
| 20978637 R | TGACACATTTGCTGCGAACAG |
| 20986398 R | TGGAAAAGATCAGTAAGGCCAGA |
| 21006046 R | ACTACATAACTTGGGTGTGTTGG |
| 21013311 R | TGCTTCTTGGTCCCGGTGT |
| ChIRP probe | |
| Probe 1 | CAGTGTTGCAGGCGAACAAA- Bio-TEG |
| Probe 2 | TTGTTCTCTGGCATAACTGG- Bio-TEG |
| Probe 3 | CACTGGAACAGTAGCTGAGA- Bio-TEG |
| Probe 4 | AGACACCATGACCAAGACGA- Bio-TEG |
| Probe 5 | CCAAAATCCACATTCTGTCA- Bio-TEG |
| Probe 6 | AACTCCTTTCAGCTTTGTTG- Bio-TEG |

| | | |
|---|---|---|
| Probe 7 | GAAGGGGCAGAGCAAGGAAA- Bio-TEG | |
| Probe 8 | GAGGCAATCTCTTGGTTTTA- Bio-TEG | |
| Antibodies | | |
| anti-LDB1 | Abcam | Cat# ab96799; RRID:AB_10679400 |
| anti-histone H3 | Abcam | Cat# ab1791; RRID:AB_302613 |
| anti-histone H3K27ac | Abcam | Cat# ab4729; RRID:AB_2118291 |
| anti-Histone H3K9me3 | Abcam | Cat# ab8898; RRID:AB_306848 |
| anti-Tubulin | Abcam | Cat# ab7291; RRID:AB_2241126 |
| anti-TBP | Abcam | Cat# ab63766; RRID:AB_1281140 |
| anti-KLF1 | Active Motif | Cat# 61233; RRID:AB_2615069 |
| anti-SET1a | Bethyl Laboratories | Cat# A700-024-T; RRID:AB_2891825 |
| anti-Pol II Ser2 | Cell signaling | Cat# 13499; RRID:AB_11378081 |
| anti-Pol II Ser5 | Cell signaling | Cat# 13523; RRID:AB_11470388 |
| anti-WDR5 | Cell signaling | Cat# 13105; RRID:AB_2620133 |
| anti-MLL1 N-Term | Cell signaling | Cat# 14689; RRID:AB_2688009 |
| anti-MLL1 C-Term | Cell signaling | Cat# 14197; RRID:AB_2688010 |
| anti-MENIN | Cell signaling | Cat# 6891; RRID:AB_10858216 |
| anti-SET1b | Cell signaling | Cat# 44922; RRID:AB_2799275 |
| anti-GAPDH | Cell signaling | Cat# 2118; RRID:AB_561053 |
| anti-Pol II | Millipore Sigma | Cat# 05-623; RRID:AB_309852 |
| anti-GATA1 | Santa Cruz | Cat# sc-1233; RRID:AB_631559 |
| anti-TAL1 | Santa Cruz | Cat# sc12984; RRID:AB_2199699 |
| anti-Pol II | Santa Cruz | Cat# sc-900; RRID:AB_2167474 |
| Normal goat IgG | Santa Cruz | Cat# sc-2028; RRID:AB_737167 |
| Normal mouse IgG | Santa Cruz | Cat# sc-2025; RRID:AB_737182 |

| | | |
|---|---|---|
| Normal rabbit IgG | Santa Cruz | Cat# sc-2027; RRID:AB_737197 |
